# Phage-encoded small RNA hijacks host replication machinery to support the phage lytic cycle

**DOI:** 10.1101/2024.10.31.621285

**Authors:** Aviezer Silverman, Raneem Nashef, Reut Bruner, Tamar Noy, Sahar Melamed

## Abstract

Bacteriophages (phages) significantly influence bacterial populations in their natural environment. However, one aspect that has not been thoroughly explored in the context of phage-bacteria interactions is the post-transcriptional regulation of gene expression, despite growing recognition of its importance in bacterial physiology over the past two decades. Important players in this process are small RNAs (sRNAs) that regulate target mRNAs via base-pairing, typically using RNA chaperones like Hfq to facilitate this regulation. Here, we apply RIL-seq to map *in-vivo* the sRNA-RNA network in *Escherichia coli* upon lambda phage infection. We highlight changes in the bacterial transcriptome and sRNA interactome while uncovering a novel phage-encoded sRNA that regulates key genes in *E. coli*. We decipher the molecular mechanism of the sRNA-mediated regulation and illustrate how the sRNA hijacks the host replication machinery and helps the infection cycle. Overall, we uncover an RNA-level regulatory layer that shapes the *E. coli* - lambda interactions.

## INTRODUCTION

Bacteriophage (phage) research has been carried out for a century now and has led and is still leading to numerous biological discoveries related to various cellular processes such as transcription, translation, replication, ligation, recombination, mutagenesis, and bacterial defense systems (e.g. CRISPR-Cas, DISARM, restriction enzymes) (reviewed in Keen et al., and Bernheim and Sorek ^1,2^). Phages are the most abundant genetic entities on Earth (estimated at 10³¹) ^3^, and they significantly impact bacterial populations and evolution (reviewed in Dion et al.^4^), contributing to bacterial virulence and antibiotic resistance ^5–7^. Phages can follow two cycles upon infection: the lytic cycle, where they replicate and lyse the bacterial cell, or the lysogenic cycle, where they integrate into the bacterial chromosome and remain inactive, replicating with the host. Virulent phages only undergo the lytic cycle, while temperate phages can enter both cycles (reviewed in Gandon ^8^).

The most extensively studied temperate phage is lambda phage. Lambda phage has a 48.5 kB dsDNA genome and was one of the first model systems for DNA research and gene expression studies (reviewed in Casjens and Hendrix ^9^). Lambda phage infects *Escherichia coli* strains, and its gene expression can be divided into immediate early, early, and late transcription. The immediate early genes, *N* and *cro*, encode proteins that manipulate host systems, primarily RNA polymerase activity and transcription termination. This manipulation allows for the expression of the early genes, followed by the late genes (reviewed in Lewis and Adhya, and Casjens and Hendrix ^9,10^). During the lytic cycle, lambda phage begins to replicate its genome through a rolling circle mechanism where the circular genome serves as a template for the synthesis of long linear molecules of lambda DNA ^11,12^. This step is followed by DNA packaging, assembly into new phage particles, and bacterial cell lysis to release new phages. (reviewed in Aksyuk and Rossmann ^13^). Depending on the sequence of events, the phage may transition to a lysogenic state, allowing it to persist longer in the host population. The lysis-lysogeny decision is made based on physiological and environmental signals, as well as the number of phages infecting each cell ^14–16^. For example, exposure to UV light, certain chemicals, and pH levels encourage the phage to excise its chromosome from the host and transition to the lytic cycle ^9,17^. The lysis-lysogeny decision involves multiple regulatory proteins encoded by the phage, mainly CI, CII, CIII, and Cro (reviewed in Casjens and Hendrix ^9^). In other temperate phages, additional factors such as short peptides can be involved in lysis-lysogeny decisions ^18,19^.

While decoding gene regulation is key to understanding the phage cycle, an aspect that was less studied in phage biology is regulation at the post-transcriptional level. In recent years, there has been increased awareness of the importance of post-transcriptional regulation of gene expression by regulatory RNAs in all three domains of life. The major group of regulatory RNAs in bacteria is comprised of short, 50-400 nt long, RNA molecules denoted small RNAs (sRNAs). Most of the sRNAs act in *trans* by base-pairing with their RNA targets upon binding the RNA chaperone Hfq and affect the target stability and/or translation (reviewed in Hör et al., and Papenfort and Melamed ^20,21^).

For many years a major unresolved challenge was the identification of the RNA pairs on Hfq. High throughput methods for transcriptome-wide identification of mRNA targets of sRNAs such as the RIL-seq (<u>R</u>NA Interaction by <u>L</u>igation and <u>seq</u>uencing) ^22,23^ and others address this need (reviewed in Wong et al., and Silverman and Melamed ^24,25^). In RIL-seq, RNAs are UV-crosslinked to an RNA-binding protein, and the protein is purified with its bound RNAs. Proximal RNA ends are ligated, yielding RNA chimeras. cDNA libraries are then built allowing the identification of individual and chimeric RNAs, which are filtered for statistically significant over-represented chimeras, generating a reliable dataset. RIL-seq was successfully applied to diverse bacterial species, including gram-negative and gram-positive bacteria, revealing thousands of novel interactions, substantial re-wiring of the network upon changes in cellular conditions, competition between RNA binding proteins, and the fact that sRNAs are encoded in all regions of the genome (reviewed in Silverman and Melamed ^24^). In general, sRNAs have been shown to play significant roles in various bacterial pathways ^21,26^, including those governed by prophage-encoded sRNAs (reviewed in Altuvia et al. ^27^).

Regulation at the RNA level of the phage life cycle was documented sporadically over the years. OOP, one of the first regulatory RNAs to be characterized ^28^, regulates the expression of CII, a positive regulator of lysogeny, in lambda phage. OOP acts as an anti-sense RNA, binding to the *cII-O* mRNA, and leading to the degradation of the mRNA ^29^. Later, prophage-encoded sRNAs that regulate targets encoded by the bacterial core genome were found (reviewed in Altuvia et al. ^27^), though very few phage-encoded regulatory RNAs were described. In the last few years, however, with the advances in RNA-seq technologies, potential phage-encoded regulatory RNAs are being detected ^30–33^. The option of cross-kingdom regulation at the RNA level between bacteria and phages was recently proposed and is an exciting route for bacteria-phage communication ^27^.

To address this option, we used the well-studied lambda phage and *E. coli* to map the RNA-RNA interaction network and the transcriptome of bacteria upon phage infection. We uncover previously under-appreciated RNA-RNA regulatory networks and document changes in the bacterial transcriptome of *E. coli* infected with lambda. We identify for the first time Hfq-dependent sRNAs encoded by lambda. Our findings highlight cross-kingdom regulation mediated by sRNAs. On the one hand, we document bacterial sRNAs that respond to phage infection, potentially serving as a defense mechanism, while on the other hand, we characterize the molecular mode of action of a phage-encoded sRNA that directly regulates essential *E. coli* genes, affecting DNA replication, and contributing to the infection process.

## RESULTS

### Profiling the transcriptome and RNA-RNA interactome under lambda infection

To monitor the changes in the *E. coli* transcriptome and identify the RNA-RNA interactome during phage infection, we grew *E. coli* MG1655 wild type (WT) carrying a chromosomal FLAG-tagged Hfq, split the cultures and infected one of the two subcultures with WT lambda phage (MOI = 5). Samples were collected at two time points, 30 min and 60 min after the infection, following exposure to UV to crosslink the RNAs and proteins. Two biological replicates were studied at each time point. Total RNA-seq and RIL-seq protocols were then applied and each fragment in a cDNA library was pair-end sequenced ^22^. As a control, cDNA libraries were similarly generated for WT cells not encoding tagged proteins (Table S1).

### Lambda phage infection affects the *E. coli* transcriptome

Total RNA-seq cDNA libraries of infected samples and uninfected samples were subjected to differential expression analyses ^34^ (Figures 1A and S1A, and Table S2), offering insights into the phage effects on the bacterial transcriptome. While the *E. coli* – lambda phage system has been studied for many years, only a limited number of studies were carried out to date to understand the effect of lambda phage or other lambdoid phages on *E. coli* by deep sequencing approaches ^35–37^. Our data reveal that the levels of several *E. coli* sRNAs, which were not known to play a role in phage infection, as well as other mRNAs, are increased upon phage infection (Figures 1A and S1A, and Table S2). Interestingly, the CpxQ sRNA, whose levels increase by ∼2-fold at 30 and 60 min following phage infection, modulates the inner membrane stress response to protect the inner membrane ^38,39^. It is reasonable to hypothesize that this upregulation of CpxQ helps the bacteria combat the phage. Together with CpxQ, CpxP levels also increase by 3-fold after 60 min but not after 30 min. CpxP is a negative regulator of the Cpx pathway and is part of the resistance to extracytoplasmic stresses ^40^, emphasizing the reaction of *E. coli* to membrane damage caused by the phage.

**Figure 1.**
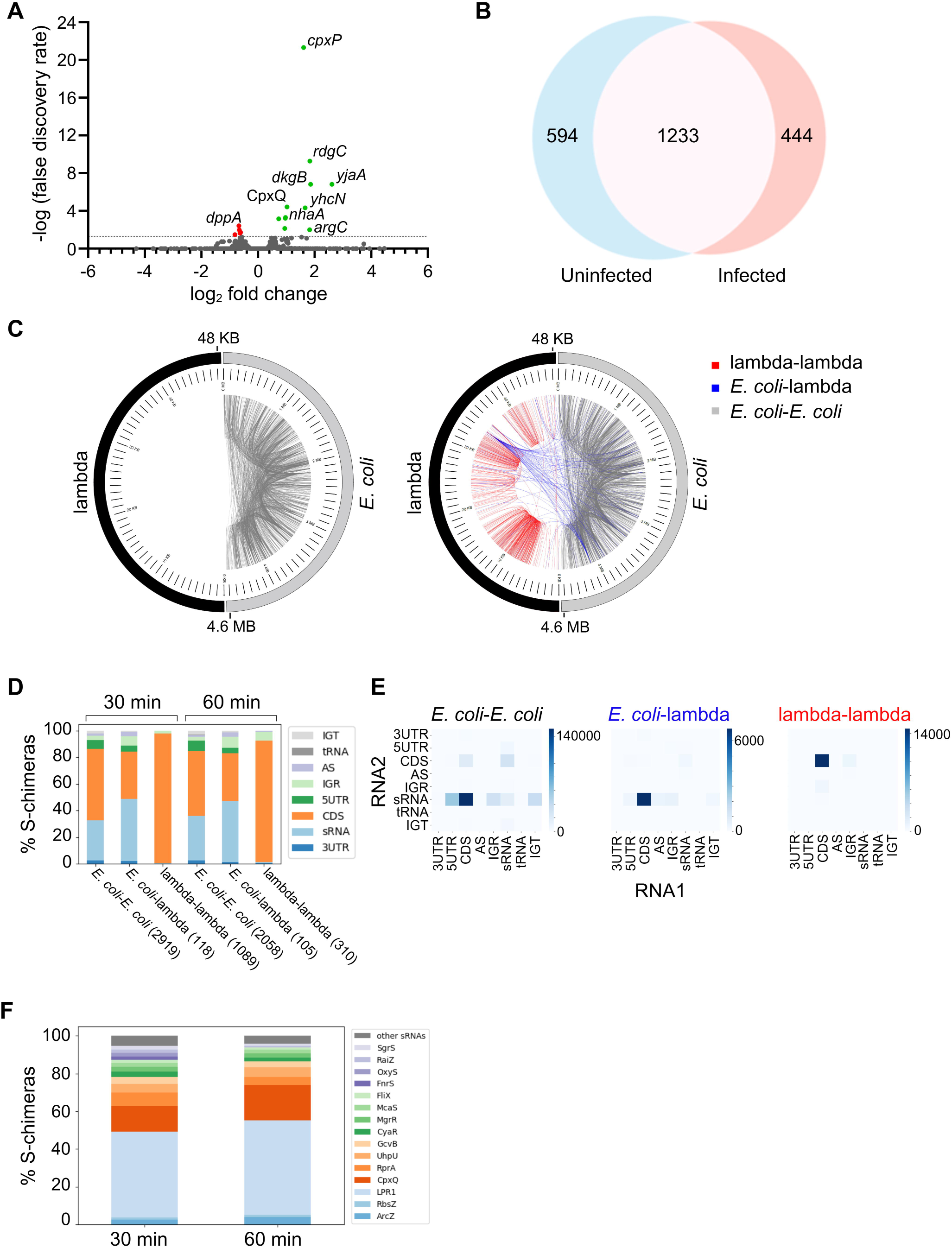
*E. coli* transcriptome and sRNA interactome during lambda infection. **(A)** Change in gene expression in *E. coli* 60 min following lambda infection is represented by a volcano plot. Genes with a p-adjusted value < 0.1 (above the dashed bar) are considered as exhibiting statistically significant changes in expression. Genes that show statistically significantly increased expression are colored green and genes that show statistically significantly decreased expression are colored red. Genes that do not statistically significantly change are colored gray. Prophage genes were removed since it was not possible to determine if the mapped reads belonged to the prophage genes or the homologous lambda genes. See more information in Table S2. **(B)** Overlap of *E. coli* chimeras between infected and uninfected bacteria. Venn diagram represents the overlap between the *E. coli* chimeras at 60 min after the infection and the chimeras obtained in *E. coli* without infection. The location of RNAs in the chimeras as RNA1 and RNA2 was ignored in this analysis. **(C)** RNA pairs of lambda–lambda and *E. coli*–lambda are detected following lambda infection. Circos plots of *E. coli* and lambda chimeras in uninfected (left) and infected samples (right) 60 min after the infection. Half of each circle represents the *E. coli* genome, and the other half represents the lambda genome. Each edge in a circle represents a chimera between two regions of the genomes. *E. coli*-*E. coli* chimeras, *E. coli*-lambda chimeras and lambda-lambda chimeras colored by gray, blue, and red, respectively. Each scale mark represents 100,000 bases in the *E. coli* half and 1,000 bases in the lambda half. **(D)** Distribution of RIL-seq S-chimeras derived from various genomic elements. Elements in S-chimeras are classified into eight categories: CDS (coding sequence), tRNA, sRNA, IGT (regions of an operonic transcript between genes), AS (antisense transcript), IGR (intergenic region), 5’ UTR, and 3’ UTR. rRNA-derived fragments were excluded. S-chimeras are separated into *E. coli*, lambda, and *E. coli*-lambda chimeras, at 30 and 60 min after infection. **(E)** Total number of chimeric fragments for each combination of genomic elements in *E. coli*-*E. coli* chimeras, *E. coli*–lambda chimeras, and lambda–lambda chimeras, 60 min after infection. Rows represent the first RNA in the chimera, and columns represent the second RNA in the chimera. **(F)** LPR1 and CpxQ dominate the *E. coli* – lambda S-chimeras that involve sRNAs. The distribution of *E. coli*–lambda S-chimeras between the different sRNAs, at 30 and 60 min after the infection.

Another intriguing transcript showing increased levels is *rdgC*. RdgC is thought to be a regulator of RecA activity by inhibiting RecA-mediated strand exchange, likely by competing with RecA for DNA binding sites ^41,42^. Increased RdgC levels may favor lambda’s lysogenic cycle by reducing RecA activity, which otherwise stimulates the autocleavage of the lambda CI repressor, a protein that maintains lysogeny ^43^. Overall, the *E. coli* transcriptome analysis upon lambda infection suggests that while most of the *E. coli* transcripts are not significantly affected by the infection, the transcripts that are affected may play a role in the relationships between the bacteria and the phage.

### RIL-seq reveals an intricate RNA-RNA network in *E. coli* during phage infection

Analysis of the sequenced fragments obtained in the RIL-seq pipeline resulted in two datasets: single fragments for which the two end sequences mapped to the same region of one genome and chimeric fragments for which the two end sequences mapped to two distinct regions within the same genome or in different genomes (e.g., *E. coli* and lambda), due to the ligation of two neighboring RNAs on the same Hfq molecule. The data reproducibility between libraries was very high, with the lowest Spearman correlation coefficient of 0.95 for single fragments and 0.76 for chimeric fragments (Figure S1B). As replicates are considered reproducible if the correlation is higher than 0.4 ^22^, the libraries were unified and the following analyses were applied to the unified datasets (Table S3). Between 2,381 to 4,126 statistically significant chimeras (S-chimeras) were obtained for each tested condition, while negligible numbers of S-chimeras were obtained for the Hfq-WT control libraries (Tables S1 and S3). These results are comparable with other RIL-seq datasets ^24^.

A comparison between infected and uninfected datasets indicates that phage infection substantially alters the sRNA-RNA interactome in *E. coli*. Specifically, 42% and 46% of the *E. coli-E. coli* S-chimeras bound to Hfq were affected at 30 min and 60 min post-infection, respectively (Figures 1B and S1C). Intriguingly, many lambda-lambda and *E. coli*-lambda RNA pairs were detected at both time points following the infection (Figures 1C and S1D). After 30 min, ∼26% of the S-chimeras involved two lambda-encoded RNAs and ∼3% were hybrids between *E. coli* and lambda phage RNAs, while after 60 min the percentages were ∼13% and ∼4%, respectively. In the *E. coli*-*E. coli* and in the *E. coli*-lambda interactions, the prominent RNA types are CDS and sRNAs while in the lambda-lambda the majority of the RNAs are annotated as CDS (Figure 1D). These data raise intriguing questions about how *E. coli-E. coli* RNA interactions are affected by phage infection, as well as the roles of both lambda-lambda and *E. coli*-lambda RNA pairs.

To this end, we analyzed the RNA pairs to identify which are the most prominent RNA pairs in the three sub-datasets (*E. coli*-*E. coli* interactions, *E. coli*-lambda interactions, and lambda-lambda interactions) based on their classification to RNA types. Among the *E. coli*-*E. coli* interactions, CDS-sRNA and 5UTR-sRNA were the most abundant type of RNA pairs, reflecting regions involved in binding between sRNAs and their mRNA targets (Figures 1E and S1E). A key player in the interactions under infection was CpxQ, which was also found in chimeras with lambda-encoded RNAs (Figure S1F). The *E. coli*-lambda RNA pairs had a similar pattern, while in the lambda-lambda interactions the most abundant RNA pairs were CDS-CDS, suggesting a different molecular mechanism for this type of interactions (Figures 1E and S1E). It is worth noting that in CDSs other RNA elements can be concealed, as was shown previously in bacterial sequences ^44^.

In these rich data of RNA-RNA interactions, we were especially intrigued to find that two RNAs were involved in 59% and 69% of the pairs that include sRNAs between *E. coli* and lambda at 30 and 60 min following the infection, respectively (Figure 1F). One RNA was the *E. coli*-encoded sRNA CpxQ and the other RNA was a distinct region in the early left operon in the lambda genome. We suspected that a sRNA is encoded in that region and termed it LPR1 (<u>L</u>ambda <u>P</u>hage <u>R</u>NA 1). LPR1 was also ranked as one of the top 15 sRNAs in chimeras forming on Hfq at both 30 and 60 min following the infection (Figure S1G), highlighting its potential relevance to Hfq-mediated regulation during lambda infection. Thus, the analysis of the RNA-RNA interaction network in *E. coli* during phage infection uncovered a mix of RNA pairs, between *E. coli-E. coli*, *E. coli*-lambda and lambda-lambda, suggesting involvement of Hfq-mediated regulation during lambda infection.

### Novel sRNAs encoded by lambda

LPR1 is located on lambda phage genome between *ea10* and *ral* and is strongly enriched on Hfq (Figure 2A). While the protein encoded by *ral* enhances restriction enzyme modification activity ^45,46^, the role of *ea10* is not fully understood. LPR1 3’ end has similar features as other Hfq-dependent sRNAs, such as a GC-rich region followed by a poly-U tail (Figure 2B). This sequence corresponds to one of the termination sites, tL2a, of lambda early left operon ^47^. To test if this region indeed encodes a sRNA, we carried out a northern analysis using RNA samples collected throughout lambda infection (Figures 2C and S2A). A top band of ∼88 nt representing LPR1 was detected in the analysis as well as a lower, stronger band of ∼68 nt. The lower band overlaps the prominent peak in the LPR1 region (Figure 2A) and represents a processing event of LPR1 to LPR1-S, which is the case for many sRNAs ^48^. LPR1 levels moderately increase from the first 30 min time point (Figure S2A) as the infection progresses (Figure 2C). Northern analysis of the RNA samples used for the RNA-seq experiment described in Table S2 showed a similar pattern, with increased levels from 30 min to 60 min (Figure S2B).

**Figure 2.**
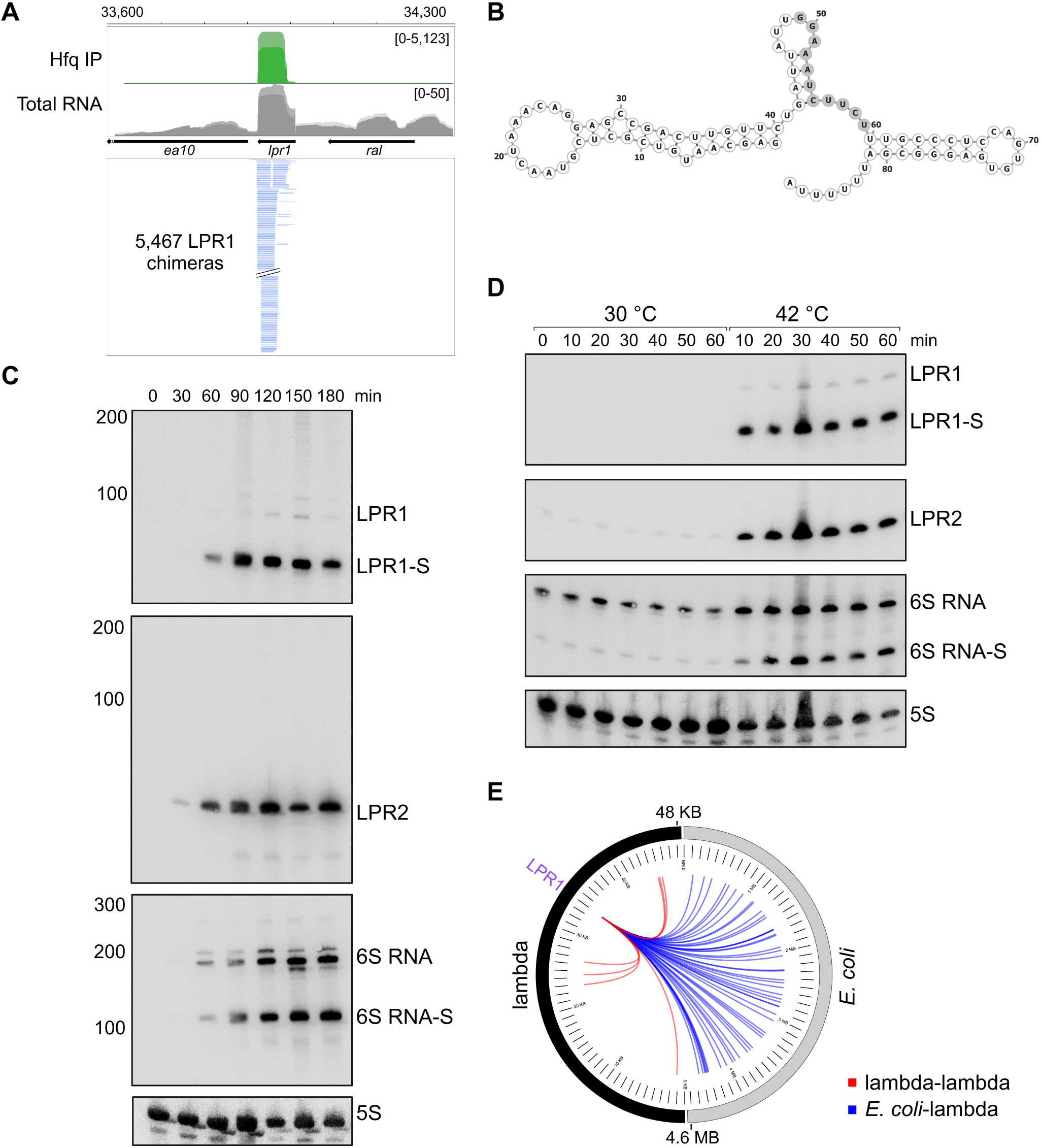
Lambda-encoded sRNAs are mainly expressed in the lytic cycle. **(A)** Browser image of *lpr1* region. Top: Hfq IP (green) and total RNA (gray). Normalized read count ranges are shown in the upper right. Bottom: chimeras of LPR1. The blue color indicates that LPR1 was found as RNA2 in the chimera. Data shown is of 30 min after the infection. **(B)** Predicted LPR1 secondary structure (drawn using Forna ^103^). Shaded bases represent the predicted seed sequence of LPR1. **(C)** Northern blot analysis detects LPR1, LPR2, and 6S RNA levels throughout the infection of *E. coli* with WT lambda phage (MOI = 0.0015). sRNAs were detected approximately 30-60 min post-infection. RNAs were probed sequentially on the same membrane, and the 5S RNA served as a loading control. Longer exposure of the LPR1 membrane is presented in Figure S2A and the same 5S RNA panel is used. **(D)** Northern blot analysis of *E. coli* carrying prophage lambda strain CI-857, grown to OD_600_ ∼0.6 at 30 °C (time point 0). Culture was split to 30 °C, representing the lysogenic cycle, and 42°C inducing the lytic cycle. LPR1 and LPR2 are detected throughout the entire lytic cycle while 6S RNA is also detected in the lysogenic state. RNAs were probed sequentially on the same membrane, and the 5S RNA served as a loading control. **(E)** LPR1 pairs with lambda-and *E. coli*-encoded RNAs. Representation of LPR1 chimeras, 30 min after infection. Circos plots were drawn as described in Figure 1C.

While investigating the RIL-seq data, we suspected that there are a couple of other novel sRNAs encoded by lambda and bound by Hfq, and the same membrane used in Figure 2C was stripped and re-probed for the two suspected regions (Figure 2C). Probing for the first region, which we termed LPR2, resulted in one defined band at the size of ∼63 nt while probing for the second region, which overlaps the previously reported lambda 6S RNA ^49,50^, resulted in two main bands, of ∼194 nt and ∼121 nt, suggesting this RNA is also processed. LPR2 and 6S RNA are found in close proximity in the lambda genome, in the late operon (Figure S2C). It is worth noting that there is no sequence similarity between the lambda 6S RNA and the bacterial global transcription regulator 6S RNA (reviewed in Cavanagh and Wassarman ^51^). Lambda 6S RNA’s role in phage biology has yet to be determined. An open reading frame, orf-64, begins within the 6S RNA and extends over the transcription terminator, when the involvement of lambda Q protein and the *E. coli* transcription elongation factor NusA cause antitermination of transcription at the terminator of 6S RNA (reviewed in Roberts et al. ^52^). This highlights the complexity of gene expression regulation in lambda.

Since lambda is a temperate phage we were curious to learn if the three sRNAs are expressed in the lytic and lysogenic cycle or only in one of them. To tease this out, we used an *E. coli* strain harboring the lambda strain CI-857, which has a temperature-sensitive CI repressor that becomes unstable at higher temperatures, enabling lytic gene expression ^53^. We grew the bacteria to mid-log phase at 30 °C to maintain lambda’s lysogenic state, split the culture, left one sub-culture at 30 °C, and shifted the other to 42 °C to induce the lytic cycle. RNA samples were collected every 10 min for 60 min from both conditions for northern analysis (Figure 2D). Strikingly, LPR1 was expressed from the first time point (10 min) during the lytic cycle, but not during the lysogenic cycle. LPR2 was also predominantly expressed in the lytic cycle, although a very faint signal could also be observed in the lysogenic state. Interestingly, 6S RNA primary transcript levels were comparable in the lytic and the lysogenic cycles, but the expression pattern of its processed transcript was different, pointing to a processing event taking place primarily during the lytic cycle. Further investigation is required to validate that it is not merely a response to the temperature shift. We were specifically interested in LPR1 that showed a strong preference for the lytic cycle and carried out another experiment in which *E. coli* was infected with a lambda strain that has an increased tendency for the lysogenic cycle, carrying CIII *tor864* ^54^, or with a lambda strain that favors the lytic cycle (CI-) (reviewed in Gottesman and Weisberg ^55^). Similar results were obtained as LPR1 was expressed under the infection with the CI-strain but not with the CIII *tor864* strain (Figure S2D). These analyses allow us to conclude that LPR1 is primarily expressed during lambda lytic cycle.

Next, we turned to study the target sets of LPR1, LPR2, and 6S RNA. Interestingly, LPR1 is always found second in the RIL-seq chimeric fragments (Figure 2A and Table S3), a feature that was found to characterize sRNAs in the RIL-seq data ^23,56^. In the RIL-seq data, LPR1 was found with dozens of targets, mainly *E. coli* encoded (Figure 2E). LPR2 and 6S RNA had substantially smaller target lists, including lambda-and *E. coli*-encoded RNAs (Figure S2E). In the current study, we decided to investigate the role LPR1 plays in the phage life cycle.

### Phage-encoded sRNA is involved in the control of spontaneous transition from lysogenic to lytic cycle

Since LPR1 was documented to interact mainly with *E. coli* targets (Figures 2E and 3A), we carried out an unbiased evaluation of this target set. We used MEME to search for a common sequence motif in the target sequences ^57^. An 11 nt sequence motif was found in all of LPR1 target sequences (Figure 3B). Examination of the sequence motif using MAST ^58^ revealed that it is complementary to the LPR1 sequence, primarily in an unpaired region (Figure 2B), suggesting a seed sequence of the sRNA, involved in base pairing to the targets.

**Figure 3.**
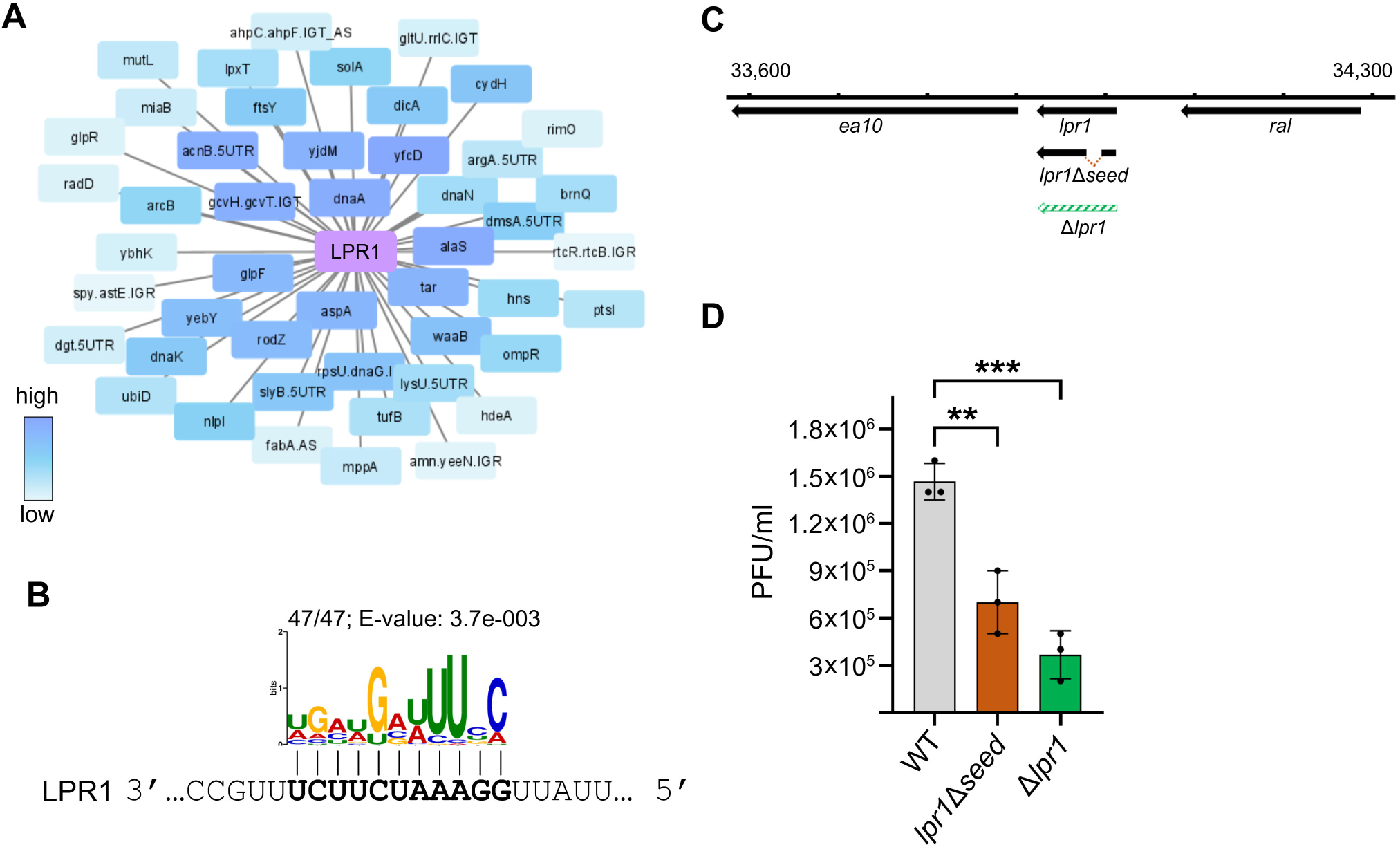
A phage-encoded sRNA is involved in the phage life cycle. **(A)** LPR1 interaction network with *E. coli* genes 60 min after infection. The color scale represents the gene’s rank based on the number of chimeric fragments it had with LPR1. The network was drawn by Cytoscape ^96^. **(B)** LPR1 target sequences share a common sequence motif that is complementary to the LPR1 sequence. Logo of the motif that was identified using MEME ^57^ in *E. coli* LPR1 target sequences 60 min after infection is shown. The fraction corresponds to the number of sequences containing the motif, over the total number of sequences analyzed. **(C)** Schematic of *lpr1* region and mutated sequences. *lpr1*Δ*seed* is colored in brown and full *lpr1* deletion is colored in green. See the Methods section for more information about the mutants. **(D)** Reduced transition of mutated prophages to the lytic life cycle. PFU/ml was detected in a spontaneous lysate of WT and mutated prophages. The two mutants show a decrease in the progeny of phages upon spontaneous induction. Results are the average of three biological replicates. Error bars represent one SD. One-way ANOVA comparison was performed to calculate the statistical significance of the change in plaque formation (**=p < 0.005, ***=p < 0.001).

To evaluate the role of LPR1 in the lambda life cycle, we deleted the entire *lpr1* sequence (Δ*lpr1*) or 18 nt that include the core of *lpr1* putative seed sequence (*lpr1*Δ*seed*) on the lambda genome (Figure 3C). Northern analysis documented comparable levels of LPR1 and LPR1Δseed (Figure S3A). We tested the effect of these mutants by monitoring the transition of lambda from the lysogenic to lytic state using a plaque assay with *E. coli* strains harboring either the WT lambda prophage or the lambda mutants. We assayed these lambda prophages in Δ*lamB* background, a gene encoding lambda phage receptor protein ^59^, to measure only the phages transitioning to a lytic state without infecting new bacterial cells (Figure 3D). Two-and four-fold reductions in the number of plaques were observed with *lpr1*Δ*seed* and Δ*lpr1* phages, respectively. In the absence of Hfq, plaque numbers for both *lpr1*Δ*seed* and Δ*lpr1* mutants were similar to that of lambda WT, supporting LPR1’s mode of action through Hfq (Figure S3B). When repeating the experiment in Δ*recA* background, impairing the autocleavage of the lambda CI repressor, a strong reduction in the number of plaques was observed between the WT *E. coli* and the Δ*recA* strains, as expected ^60^ (Figure S3C). However, no difference in the plaque number was observed between the WT lambda and its mutants in the Δ*recA* background, hinting that LPR1 activity is downstream and dependent on the autocleavage of CI. Together, these results suggest a role for LPR1 in the transition to or the support of lambda lytic cycle.

### LPR1 directly regulates essential *E. coli* genes and affects phage DNA replication

At 60 min post-infection, RIL-seq identified 47 targets of LPR1, nine of which were essential *E. coli* genes (*ftsY*, *ptsI*, *alaS*, *dnaN*, *rpsU*, *dnaG*, *dicA*, *dnaA*, *ubiD*). Out of the 4,720 annotated *E. coli* genes, 358 are essential ^61^. A Fisher’s exact test revealed that the nine essential genes found in chimeras with LPR1 were statistically significantly enriched (p-value = 0.007), giving the general abundance of essential genes in *E. coli*. This finding suggests that LPR1 preferentially binds to critical, essential genes in the host rather than associating randomly with other genes ^62^.

We decided to test the regulation by LPR1 of the two essential target genes with the highest number of chimeric fragments, *dnaA-dnaN* region (termed *dnaAN*) and *alaS*. *dnaA* and *dnaN* are essential genes encoded in an operon that includes *recF*. These two genes encode components of the *E. coli* DNA replication machinery; DnaA is a replication initiator protein (reviewed in Hansen and Atlung ^63^) and DnaN is β sliding clamp that links the DNA polymerase to the DNA ^64^. AlaS is a tRNA-ligase that plays a part in the incorporation of alanine during the translation process ^65^. It is an essential gene in *E. coli* and is autoregulated. We fused a region of *dnaAN* or *alaS*, including the position of chimeras with LPR1 (Figures 4A and S3D), and the region of predicted base-pairing (Figures 4B and S3E), to a GFP reporter, creating a translational fusion ^66,67^. In parallel, we cloned LPR1 under an IPTG-induced promoter in the pNM46 plasmid ^68^ (Figure S3F). LPR1 overexpression elevated the expression of the *dnaAN*-gfp fusion by more than 4-fold, indicating a positive regulation of *dnaAN* by LPR1 (Figures 4C and D). A 3 nt mutation in the LPR1 predicted base-pairing sequence (LPR1-M1) eliminated the up-regulation, while a compensatory mutation on *dnaAN* sequence restored the ∼4-fold up-regulation (Figure 4D). These results provide support for the direct interaction between LPR1 and *dnaAN* and for the regions involved in base pairing. On the contrary, LPR1 overexpression resulted in a ∼2-fold reduction of the GFP signal for the *alaS-gfp* fusion, suggesting versatile modes of action for LPR1 (Figure S3G). A 3 nt mutation in the LPR1 predicted base-pairing sequence (LPR1-M2) eliminated the down-regulation, while a compensatory mutation on *alaS* sequence restored the ∼2-fold down-regulation (Figure S3G).

**Figure 4.**
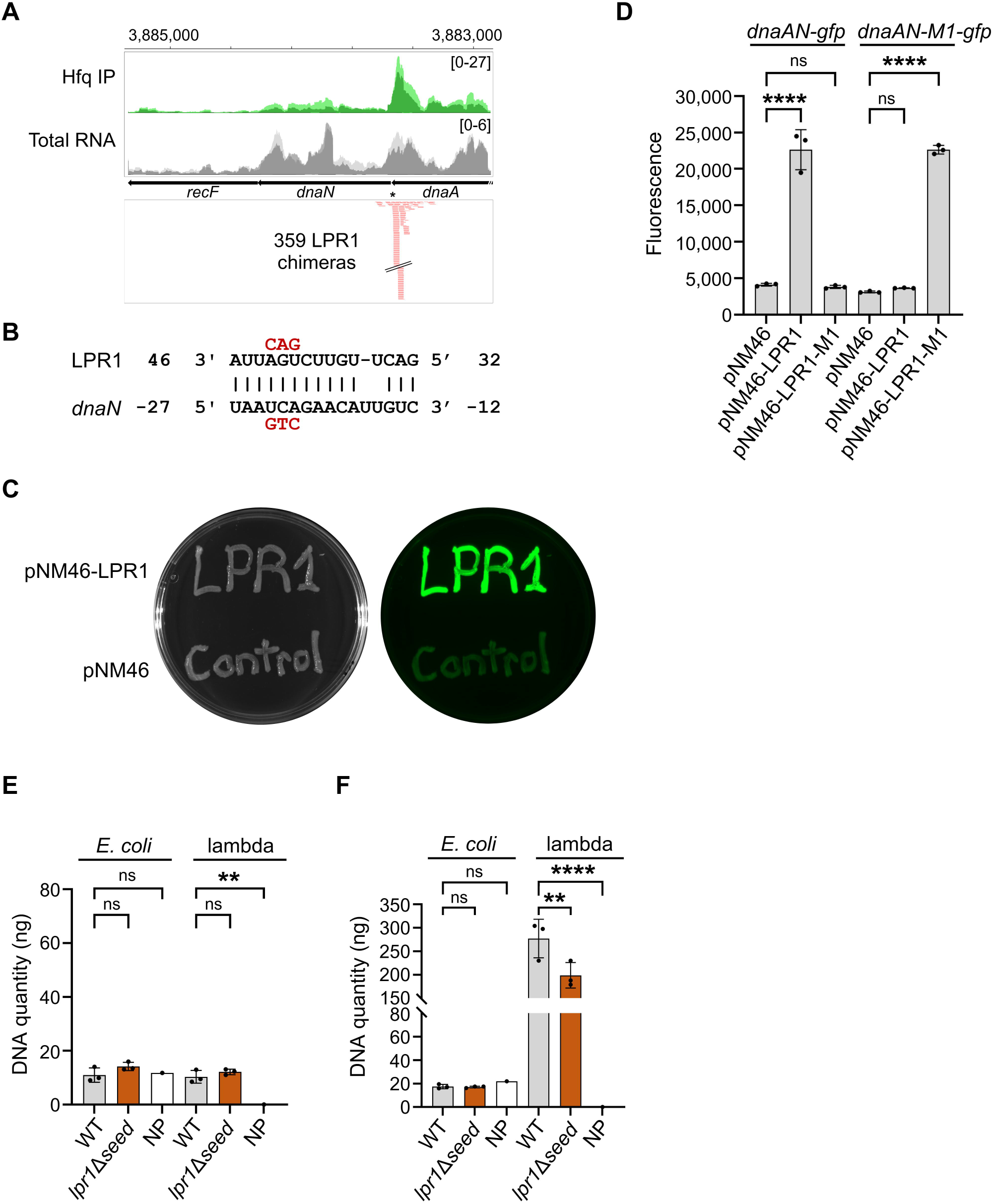
LPR1 upregulates *dnaN* and plays a role in phage DNA replication. **(A)** Browser image of *dnaAN-recF* region. Top: Hfq IP (green) and total RNA (gray). Normalized read count ranges are shown in the upper right. Bottom: chimeras of *dnaAN* with LPR1. The red color indicates that *dnaAN* was found as RNA1 in the chimera. The asterisk indicates the position of the base pairing with LPR1. Data shown is of 30 min after the infection. **(B)** Base pairing between *dnaN* and LPR1 with sequences of mutants assayed (colored in red). Numbering is from AUG of *dnaN* mRNA and +1 of LPR1 sRNA. **(C)** LPR1 overexpression increases *dnaAN-gfp* signal. Bacteria expressing *dnaAN-gfp* from pXG10-SF carrying a control vector (pNM46; bottom) or overexpressing LPR1 (pNM46-LPR1; top), were plated on agar plates supplemented with 1 mM IPTG. White light (left) and fluorescent signal (right) images were taken using ChemiDoc imager. **(D)** LPR1 induces the expression of *dnaAN-gfp* reporter fusion using *dnaAN-gfp* expressed from pXG10-SF. However, this activation is not observed when using LPR1-M1. Compensatory mutation in *dnaAN-gfp* restores the upregulation by LPR1. Average fluorescent values are based on three biological replicates. Error bars represent one SD. One-way ANOVA comparison was performed to calculate the statistical significance of the change in GFP signal (ns = not significant, ****=p < 0.0001). **(E-F)** lambda DNA levels are lower in *lpr1*Δ*seed* in comparison to WT lambda 60 min following induction of prophages by UV as measured by qPCR. DNA levels before UV irradiation are comparable for *E. coli* and lambda in both of the prophages. One-way ANOVA comparison was performed to calculate the statistical significance of the change in DNA levels (ns = not significant, **=p < 0.005 ****=p < 0.0001).

We hypothesized that LPR1 positively regulates components of the DNA replication machinery to benefit its own genome replication. To test the connection between the regulation of *dnaAN-gfp* fusion and the effect on phage DNA replication we measured the levels of *E. coli* and lambda DNA in *E. coli* harboring WT lambda or *lpr1*Δ*seed*. Cultures were grown to mid-log phase, exposed to UV irradiation, and then returned to the incubator. Samples were collected before irradiation and 60 min after to quantify *E. coli* and lambda DNA levels by qPCR (Figures 4E and 4F). In their lysogenic state, both strains had similar levels of lambda DNA and *E. coli* DNA, reflecting the incorporation of the phages into the bacterial chromosome. Following lambda lytic cycle induction by UV irradiation, lambda DNA levels increased dramatically versus *E. coli* DNA in the WT lambda background, but to a lower extent in *lpr1*Δ*seed* background, supporting a positive role of LPR1 in lambda DNA replication.

### LPR1 enhances *dnaN* translation initiation

Next, we aimed to determine the molecular mechanism underlying the positive regulation of *dnaAN-gfp* fusion by LPR1. In our RNA-seq data (Table S2), there was no significant change in the levels of *dnaA*, *dnaN,* or *recF* upon lambda infection. This led us to hypothesize that LPR1’s regulation of this target is at the translational level rather than on RNA stability. We carried out a western analysis in which each of the genes in the *dnaAN-recF* operon was tagged with SPA in the presence of a control vector, LPR1 overexpression, or LPR1-M1 overexpression. DnaA-SPA and RecF-SPA levels were not affected by the presence of LPR1 or LPR1-M1 whereas DnaN-SPA levels increased in the presence of LPR1 but not in the presence of LPR1-M1 (Figure 5A). These results suggest that LPR1 specifically targets *dnaN* and not the entire operon.

**Figure 5.**
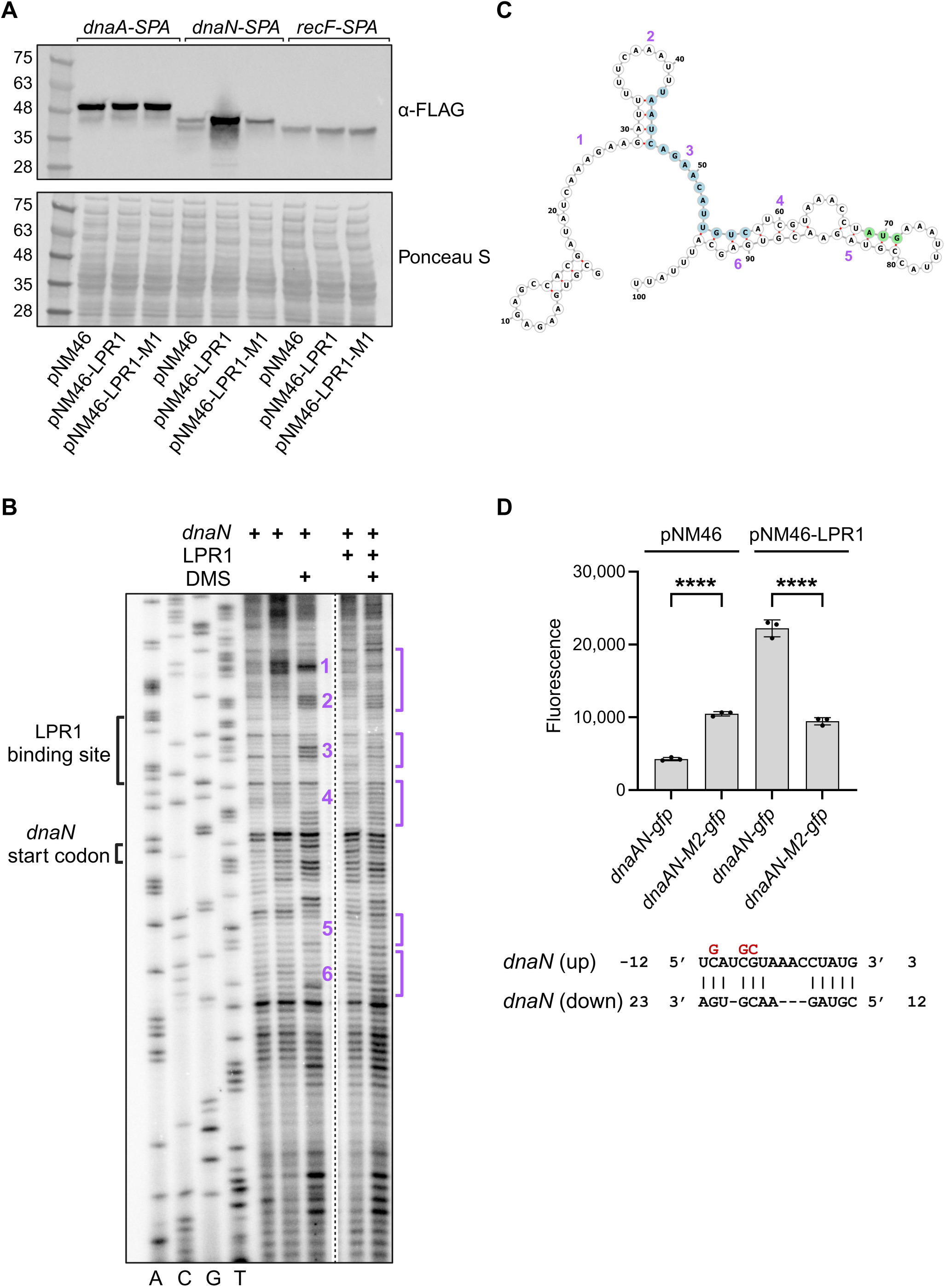
LPR1 exposes the *dnaN* RBS to enhance translation initiation. **(A)** immunoblot analysis showing that LPR1 upregulates DnaN-SPA levels but not DnaA-SPA or RecF-SPA. The three genes in the operon were tagged with an SPA tag and their levels were tested with pNM46, pNM46-LPR1, or pNM46-LPR1-M1 by immunoblot analysis using α-FLAG antibody. Ponceau S served as a loading control. **(B)** Structural probing of *dnaN* with and without LPR1 treated with DMS supports direct base-pairing between the two RNAs. DMS methylates unpaired adenine and cytosine. Regions where binding of LPR1 changes the DMS methylation pattern are numbered (1-6), while region 3 overlaps the LPR1 binding site as shown in Figure 4B. The left four lanes are Sanger sequencing of the *dnaN* sequence used in this assay, ACTG represents the ddNTP that was added. Note that the observed band pattern represents the *dnaN* complementary sequence. Lanes 5 and 6 are *dnaN* without-and with -being heated to 70 °C, respectively. Lane 7 is *dnaN* treated with DMS. Lanes 8 and 9 are *dnaN* treated with DMS without – and with – LPR1, respectively. In all lanes, a radioactive primer (P1102) was used for the primer extension. **(C)** Predicted secondary structure of the *dnaN* RNA used for the structural probing assay (drawn using forna ^103^). The first 100 nt of the *dnaN* RNA used are shown. LPR1 binding site is highlighted in blue, *dnaN* start codon is highlighted in green, and the positions changed in the structural probing assay are labeled with purple numbers. **(D)** Mutation upstream to *dnaN* CDS interrupts the secondary structure involving *dnaN* RBS. The *dnaAN-M2-gfp* fusion signal is elevated in comparison to the *dnaAN-gfp* signal, recapitulating the effect of LPR1 on the secondary structure. *dnaN-gfp* reporter fusion induction by LPR1 is eliminated in *dnaN-M2-gfp*. Average fluorescent values are based on three biological replicates. Error bars represent one SD. One-way ANOVA comparison was performed to calculate the statistical significance of the change in the GFP signal (****=p < 0.0001).

Since LPR1 base pairs with *dnaN* upstream to its start codon (Figures 4A and 4B), we hypothesized that LPR1 increases the production of the DnaN protein by increasing its translation initiation rate by exposing the *dnaN* ribosome binding site (RBS). To test this, we monitored the effect of LPR1 on the *dnaN* mRNA secondary structure by in-vitro structural probing in the presence or the absence of LPR1 under Dimethyl sulfide (DMS) treatment or RNase T1 digestion (Figures 5B and S4A). We noticed several distinct changes in *dnaN* mRNA secondary structure in the presence of LPR1. These changes were mainly positioned upstream to the *dnaN* start codon, overlapping the base-pairing region of LPR1 and the *dnaN* RBS (Figures 5C and S4B). With DMS treatment, in regions 1-3, a reduction in band intensity is observed following LPR1 binding, suggesting a transition from ssRNA to dsRNA formation. In regions 4-6, which overlap with the *dnaN* RBS and its CDS, an increase in band intensity is observed, suggesting a transition from dsRNA to ssRNA (Figure 5C). A similar trend is observed with RNase T1 digestion (Figure S4A). These changes support our hypothesis, that the *dnaN* RBS is exposed upon binding of LPR1, enabling increased translation initiation of *dnaN*. We also mutated the *dnaAN-gfp* fusion sequence (*dnaAN-M2-gfp*) upstream to the *dnaN* start codon ((-10)-(-8) nt), disrupting the base-pairing of that region with downstream *dnaN* sequence, exposing the RBS. With just the control vector, the GFP signal of the mutated *dnaAN-gfp* increased by ∼2-3 fold in the absence of LPR1, supporting a stronger translation initiation due to the introduction of the mutation (Figure 5D). LPR1 upregulation of the *gfp* fusion was completely eliminated in *dnaAN-M2-gfp*, likely because the *dnaN* RBS was already exposed in the mutant. This further supports our model that LPR1 is upregulating *dnaN* expression by affecting its translation initiation.

### LPR1 is conserved across diverse phage and bacterial sequences

To evaluate the prevalence of LPR1 in the phylogenetic tree, we performed a search for sequences similar to LPR1 using BLAST and NCBI nucleotide database (Figure 6A). LPR1 sequence was found to be conserved (with an identity of at least 92%, and with no more than three gaps) in other lambdoid phages and another 8 phage species. In addition, LPR1 was found to be conserved in 8 different bacterial species, likely as part of a prophage. In many of these species, conservation extends to LPR1’s adjacent genes, suggesting that these regions may work together in important, coordinated ways (Figures 6B and S5B). Altogether, these results showing LPR1 conservation in different phages and bacteria suggest it may play a broader role in the interactions between bacteria and phages.

**Figure 6.**
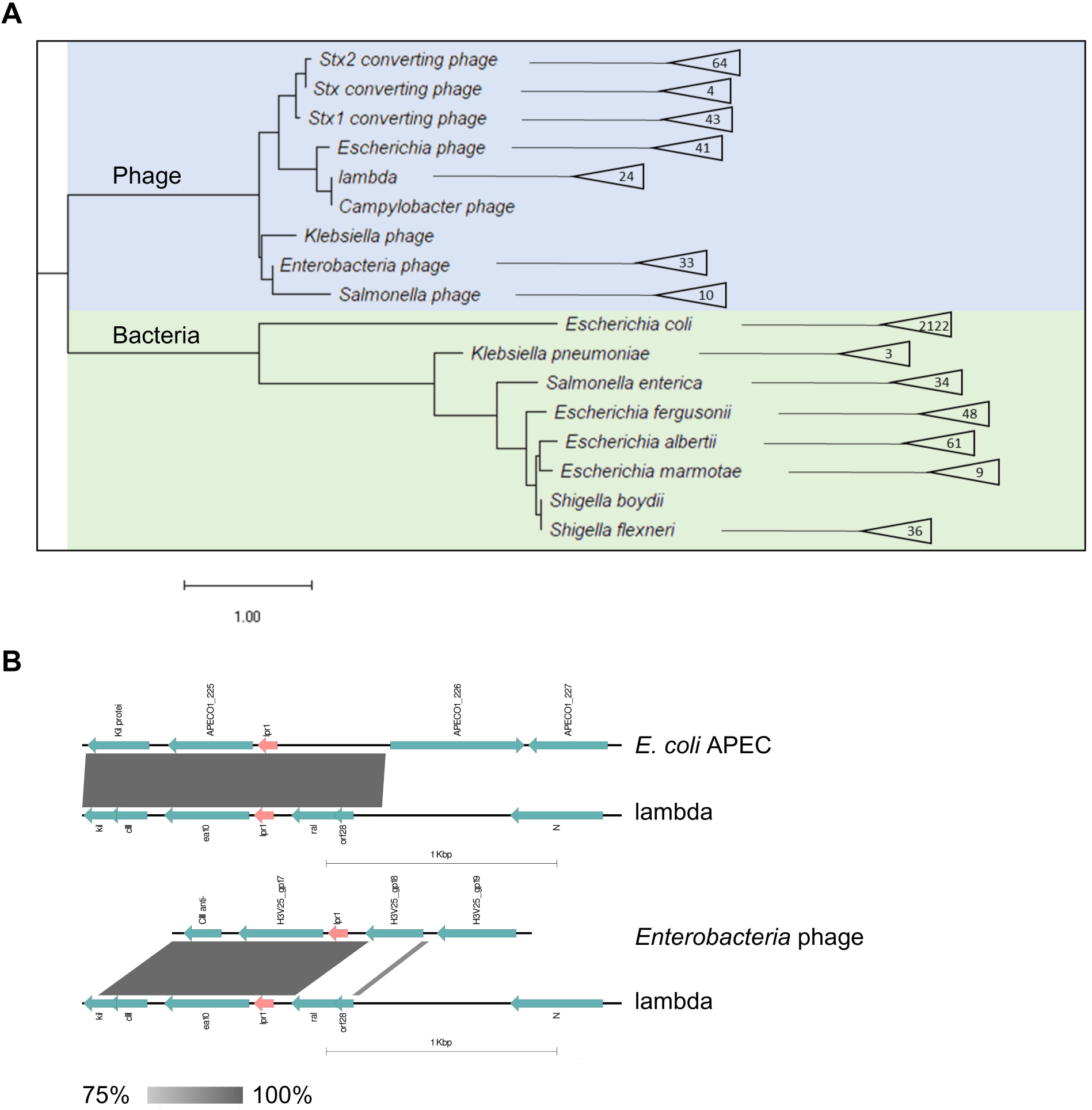
Conservation of LPR1 across bacteria and phages. **(A)** LPR1 is conserved across many bacterial and phage sequences. A phylogenetic tree of the genomes that LPR1 was conserved in, according to BLAST results. LPR1 sequence was found in 2,537 relevant genomes from the NCBI nucleotide DB. The tree was generated according to one representative genome for each species (Table S4). The green tree represents bacterial species while the blue tree represents phage species. The numbers in the triangles are the number of genomes in which LPR1 was found for each species. A scale is shown at the bottom, where the distance between the bacterial species and the phage trees is arbitrary. **(B)** Conservation between lambda and *E. coli* APEC or Enterobacteria phage in the region of LPR1. The similarity between regions on the genomes is marked by gray. The scale represents the identity percentage according to BLAST. The full names of the sequences are listed in Table S4.

### Model of LPR1 (PreS) mode of action

Collectively, our results suggest that LPR1 supports lambda lytic cycle through upregulation of *dnaN* and we decided to rename it as PreS (<u>P</u>hage replication <u>e</u>nhancer <u>s</u>RNA). Next, we were intrigued to learn if PreS’s mode of action on *dnaN* could be predicted using 3D modeling software. We turned to AlphaFold 3 ^69^ to compare our experimental results of the Hfq*-dnaN*-PreS interaction with the software 3D model prediction (Figures 7A, S6A-B, and Supplemental Video 1). Nicely, the 3D model of the Hfq*-dnaN*-PreS interaction recapitulated our experimental results (Figures 5 and S4) and the known principles of sRNA-mRNA interactions taking place on Hfq. PreS’s (labeled in green) poly-U tail is embedded in the Hfq hexamer (labeled in pink) proximal side and *dnaN* (labeled in turquoise) is bound to the distal side of the Hfq hexamer, as was shown for other sRNA-mRNA pairs ^70^. According to the 3D model, the binding regions of PreS and *dnaN* are found in close proximity to each other, allowing base pairing, while the RBS (labeled in gray) does not form a strong secondary structure and is available for ribosome binding.

**Figure 7.**
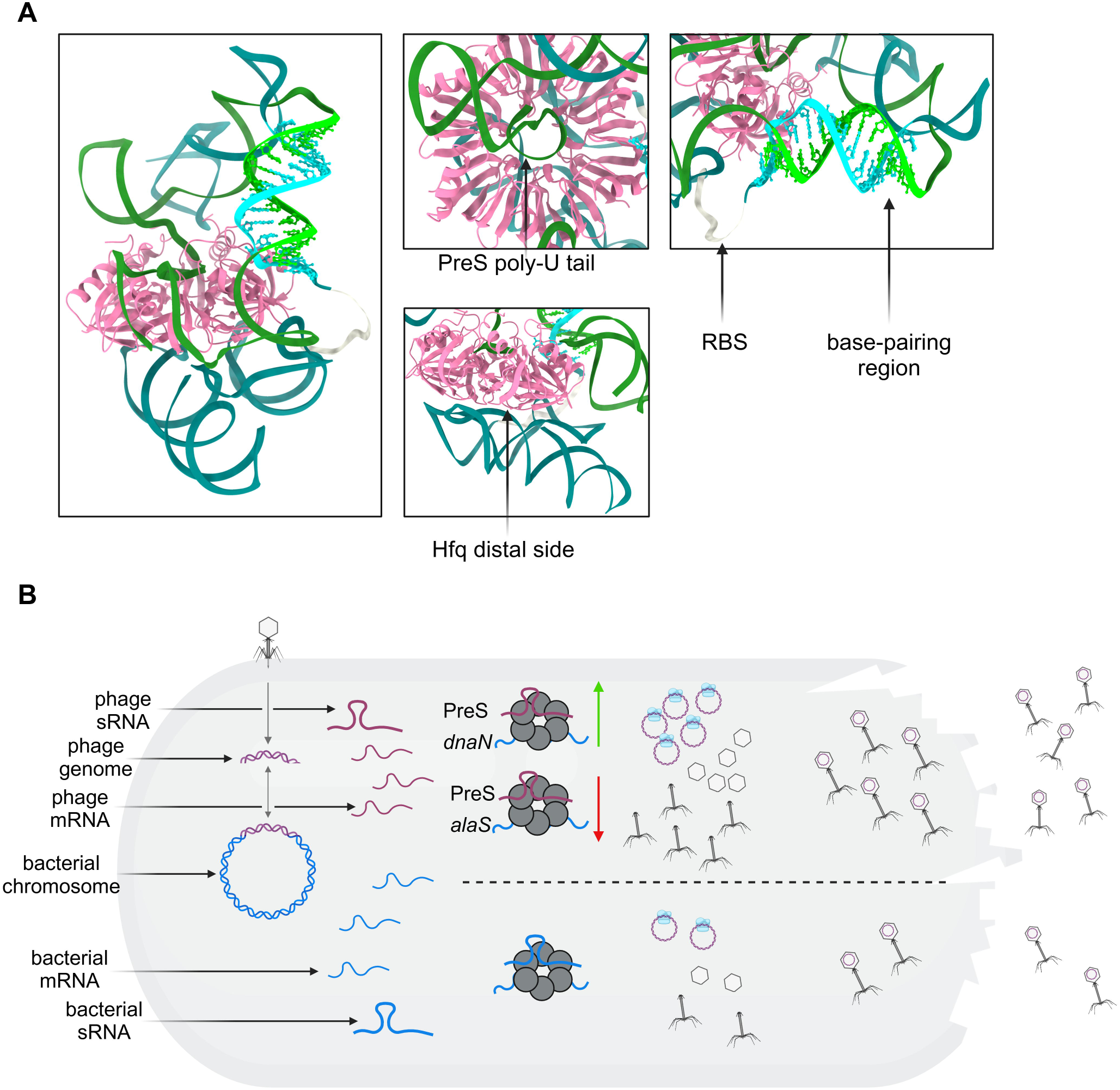
Model for LPR1 (PreS) mode of action. **(A)** 3D model of the Hfq*-dnaN*-PreS complex. Prediction of 3D structure of the Hfq*-dnaN-* PreS complex by AlphaFold 3 ^69^ recapitulates our experimental results for *dnaN*-PreS interaction. Hfq hexamer is colored pink. *dnaN* is colored turquoise while its RBS is colored gray. PreS is colored green. **(B)** When the lambda genome is incorporated into the bacterial chromosome, it can be excised spontaneously or by environmental stresses. When phage transcripts are being transcribed, some are being bound by Hfq, such as PreS (above the dashed bar). PreS interacts with *E. coli-*encoded transcripts on Hfq and regulates their expression, either positively (e.g. *dnaN*), or negatively (e.g. *alaS*). PreS-mediated regulation assists lambda DNA replication (DNA polymerase is colored in light blue) and the lytic infection cycle. When PreS is absent (below the dashed bar), lambda replication decreases, and fewer new phage particles emerge.

We suggest the following model for the mode of action of PreS (Figure 7B). The integration of the lambda genome into the bacterial chromosome allows for spontaneous excision or excision induced by environmental stresses. This process triggers the transcription of phage RNAs, including PreS, which interacts with Hfq. Through this interaction, PreS modulates the expression of *E. coli*-encoded transcripts, regulating them both positively (e.g., *dnaN*) and negatively (e.g., *alaS*). The positive regulation of *dnaN* by PreS relies on increase in translation initiation. This regulatory activity of PreS is instrumental in supporting lambda DNA replication and facilitating the lytic infection cycle. It is worth noting that the RIL-seq analysis captured many other lambda and *E. coli* RNA pairs on Hfq, which are expected to contribute to our understanding of phage-bacteria interactions.

## DISCUSSION

In this study, we mapped the sRNA interactome of *E. coli* during lambda phage infection, expanding the knowledge of sRNA networks to host-phage relationships. While providing thorough analyses of the changes in the bacteria and phage transcriptomes, and sRNA interactome, we discovered previously overlooked lambda-encoded sRNAs. The discovery of novel sRNAs encoded by lambda phage that rely on the bacterial Hfq protein provides new insights into how phages can manipulate host regulatory networks, revealing potential mechanisms through which phage sRNAs may influence host gene expression and enhance the efficiency of the phage life cycle. These sRNAs represent a previously unrecognized layer of regulatory complexity that phages utilize to manipulate their bacterial hosts.

### Accessibility of transcriptome and RNA-RNA interactome data

In the current work, as is the case in many of the studies carried out these years, valuable datasets are generated by RNA-seq-based methodologies. To make our current RNA-seq and RIL-seq data accessible and user-friendly, the data were uploaded to the UCSC Genome Browser ^71^ and can be visualized at: (*link will become available upon publication*). Visualization of the RNA-seq and RIL-seq data can assist in the identification of new sRNAs, assessment of gene expression patterns, discovery of new interactions, and potentially new biological concepts.

### Versatility of sRNA mode of action

Bacterial sRNAs exhibit remarkable versatility in their modes of action, enabling the regulation of a wide range of cellular processes. Unlike protein-based regulatory mechanisms, base-pairing with target mRNAs leads to diverse outcomes such as mRNA degradation, stabilization, or translational inhibition/activation. This RNA-mediated regulation allows for a rapid and dynamic response to environmental changes. Our findings in this study highlight how the versatility of sRNAs extends to phage-encoded sRNAs. PreS acts as a repressor when it downregulates *alaS* expression, while at the same time, it acts as an activator when it upregulates *dnaN* expression. The use of the same molecule for performing both of these types of regulation benefits the phage, allowing it to multiply more efficiently. Our data suggests that PreS has dozens of other targets besides *dnaN* and *alaS*, examination of which would help us understand the extent to which the versatility of PreS reaches.

### Role of phage-encoded sRNAs in phage biology

The interactions between viruses and their hosts are complex and involve diverse factors on multiple levels. Upon infection, lambda phage exploits various host systems to facilitate its progression toward either the lytic or lysogenic cycle ^72^. Similarly, in other phages, it has been shown that they produce regulatory molecules to sabotage bacterial transcription machinery ^73^. In eukaryotes, viral miRNAs play a critical role in modulating host gene expression through the targeted degradation or translational repression of specific mRNAs, often utilizing RNA chaperones such as Argonaute ^74–77^. Here, we demonstrate the involvement of Hfq-dependent sRNAs, which were generally overlooked for their contribution to the arms race between bacteria and phages at the post-transcriptional level. We discover Hfq-dependent sRNAs encoded by lambda phage that contribute to the phage life cycle.

One of these sRNAs, PreS, is involved in fine-tuning the expression of essential bacterial genes during phage infection and represents an adaptive mechanism evolved by lambda phage to enhance its survival and replication efficiency. By targeting the *dnaN* host gene, which is involved in DNA replication and is thus critical for the phage life cycle, PreS helps lambda optimize the cellular environment for phage replication. Interestingly, *dnaN* expression is upregulated in an SOS-dependent manner following UV irradiation and exposure to alkylating agents ^78,79^. Under these stress conditions, DnaN is involved in mediating the recruitment of error-prone polymerases for translesion synthesis and initiating methyl-directed mismatch repair in response to the stress. Furthermore, it was previously suggested that there is post-transcriptional activation of *dnaN* upon UV or nalidixic acid treatment but no mechanism was reported ^80^.

Herein, we provide a model by which *dnaN* is activated at the post-transcriptional level by exposure of its RBS (Figure 5). UV irradiation and exposure to DNA alkylating agents are also the triggers for the transition of lambda phage from a lysogenic to a lytic life cycle. This information supports the model that PreS plays a role in the lytic cycle of lambda phage, potentially influencing the switch between the lytic and lysogenic pathways. Future work, focusing on PreS lambda-encoded targets, could further clarify this model.

While we deciphered the significance of *dnaN* upregulation for lambda phage, the significance of downregulating *alaS* still needs to be uncovered. By limiting the availability of charged tRNA for alanine, the phage may inhibit the host protein synthesis, reallocating cellular resources toward its own replication and viral protein production. Additionally, this suppression could trigger stress responses, such as the stringent response ^81^, which halts normal bacterial growth or interferes with peptidoglycan synthesis that contains alanine.

The two other Hfq-dependent sRNAs found in our study, LPR2 and 6S RNA, are encoded in the late operon and may play different roles in lambda biology. Interestingly, LPR2 was found only with two other RNAs in the RIL-seq data, PreS and the *E. coli*-encoded *fliC*. In previous studies, sRNAs that had a very limited number of targets were often found to act as RNA sponges ^23,56^. The possibility of LPR2, encoded in the late operon, to sponge PreS, encoded in the early left operon, is an intriguing possibility for complex post-transcriptional regulation between lambda transcripts. 6S RNA has several *E. coli*-encoded targets and one lambda early operon-encoded target. Interestingly, a 74-nt sRNA named StxS, characterized in Shiga toxin-encoding phages ^82^, was found in the same region of LPR2 and 6S RNA. However, no sequence similarity was detected between StxS and these sRNAs. StxS was found to repress Shiga toxin 1 expression under lysogenic conditions by interacting with the stx1AB transcript and to activate the general stress response factor of *E. coli*, RpoS, through interactions with its 5’ UTR. This activation supports cell growth under nutrient-limited conditions. Further work is needed to determine the role LPR2 and 6S RNA play in phage-host interactions.

### Prevalence of phage-encoded sRNAs

The interaction between lambda phage and *E. coli* is a subject of significant research interest due to its implications for both phage biology and host defense mechanisms. The identification of Hfq-dependent sRNAs encoded by lambda phage opens up new perspectives towards the molecular mechanisms of phage-host interactions. This discovery suggests a sophisticated strategy by which phages can manipulate host cellular processes to their advantage. We were excited to find that PreS is highly conserved in hundreds of sequences beyond *E. coli* (Figures 6 and 5S). The broad distribution of PreS across phylogenetic groups aligns with recent findings from the Papenfort lab, which identified a lysogenic phage-encoded sRNA, VpdS, that regulates the prophage-host interaction in *Vibrio cholerae* ^83^. The phage is induced by a quorum sensing signal and when VpdS is being produced, it regulates host and phage mRNAs. The parallel evolution of the two sRNAs, PreS and VpdS, both contributing to the phage lytic cycle, suggests that the role of sRNAs in phage biology may be more fundamental than previously recognized. Further investigation into the role of these sRNAs and others could provide valuable insights into the dynamics of phage infections and the evolution of phage strategies for host manipulation.

Given the renewed interest in phage biology over the past decade ^84^, it is anticipated that additional phage-encoded sRNAs will be identified, uncovering novel mechanisms by which phages hijack and manipulate host cellular machinery for their own advantage. Growing resources of phage banks ^85,86^ and other phage collections ^87^ are invaluable resources for conducting such studies and further elucidating the role of sRNA-mediated regulation on a broader scale.

## METHODS

### LEAD CONTACT AND MATERIALS AVAILABILITY

Further information and requests for resources and reagents should be directed to and will be fulfilled by the Lead Contact, Sahar Melamed.

### EXPERIMENTAL MODEL AND SUBJECT DETAILS

#### Bacteria

*E. coli* MG1655 (crl^+^ or crl^-^) strains served as the WT strains in this study. All other bacterial strains are listed in Table S5 along with plasmids and oligonucleotides used. Unless indicated otherwise, all strains were grown with shaking at 250 rpm at 37 °C in an LB-rich medium or TBMM. Ampicillin (100 µg/ml), chloramphenicol (30 µg/ml), kanamycin (30 µg/ml), arabinose (10 mM or 0.2%), and IPTG (1 mM) were added where appropriate. Unless indicated otherwise, overnight cultures were diluted to OD_600_ = 0.05 and grown for the indicated times or to the desired optical densities. MG1655 prophage was constructed by recovering a colony from the center of a turbid plaque. The incorporation of the lambda genome into the bacterial chromosome was verified by colony PCR using primers P810 and P811 (Table S5).

#### Phages

Lambda phage WT was kindly provided by Susan Gottesman. To test specific life cycles, the following mutants were used: (CI-) - obligatory lytic, (CIII864) - favoring the lysogenic cycle, and (CI857) - a temperature-sensitive CI mutant. These mutants were kindly provided by Shoshy Altuvia and Sivan Pearl-Mizrahi as described in the Acknowledgements section. Unless indicated otherwise, all phages were grown with their host in TBMM: Tryptone, NaCl, and 20mM MgSO_4_, and supplemented with maltose (0.2%) and vitamin B1 (0.0001%). When appropriate, phages were diluted in TM buffer (1 M Tris 7.5ph and 10 mM MgSO_4_). The induction of prophages was done by 40 mJ/cm^2^ of 254 nm UV irradiation using UVP Crosslinker (Analytik Jena).

### METHOD DETAILS

#### Plaque assay and determination of multiplication of infection (MOI)

WT *E. coli* was grown in TBMM supplemented with maltose and vitamin B1 to OD_600_ of 0.4-0.6, then 100 μl of serial dilutions from stock phage were mixed with 100 μl of bacteria. After incubation for 20 min at 37 °C with rotations, 3 ml of top agar (TBMM 0.7% agar) was added to the bacteria and the phages, vortexed, and immediately plated on LB agar plates (1.5% agar). Following solidification, plates were incubated overnight at 37 °C. Plaques were counted and PFU\ml was calculated. CFU\ml of the *E. coli* host strain was measured by plating serial dilutions of bacteria throughout its growth curve. MOI was calculated as the ratio between PFU\ml and CFU\ml.

#### Plasmid construction and mutagenesis

*E. coli* K-12 MG1655 genomic DNA was used as a template to amplify mRNAs to be cloned into the respective constructs. Cloning LPR1 into pNM46, a pBR carrying the *lacI* gene and an IPTG inducible promoter ^68^, was done by amplifying the vector using primers P814 and P815 (Table S5). LPR1 was amplified by PCR from purified prophage DNA using primers P810 and P811 (Table S5) which had flanking sequences homologous to the insert site on pNM46. PCR products were ligated according to the NEBuilder assembly master mix protocol (New England Biolabs). Target sequences were cloned into pXG10-SF plasmid as follows. Regions of target genes, mainly regions captured in the chimeric fragments, were PCR amplified, digested with Mph1103I and NheI, and cloned into pXG10-SF digested with the same restriction enzymes. The inserted sequences were verified by colony PCR and Sanger sequencing.

Mutagenesis of the different plasmids was carried-out using the QuikChange Lightning Site-Directed Mutagenesis Kit (Agilent). All plasmids were freshly transformed into the appropriate strains before each experiment.

#### RNA isolation

Cells corresponding to the equivalent of OD_600_ between 5 to 10 were pelleted, washed once with 1 X PBS, and frozen in liquid nitrogen. RNA was extracted according to the standard TRI Reagent protocol (Sigma-Aldrich) as described previously ^56^. At the final step, RNA was resuspended in 20–50 µl of DEPC-treated water and quantified using a NanoDrop (Thermo Fisher Scientific).

#### Northern blot

Total RNA (10 μg) was mixed with 1 X RNA loading dye (Thermo Fisher Scientific). RNA was heated at 90 °C for 4 min, and loaded onto a denaturing 6% Acrylamide urea gel containing 8 M urea, in 1 X TBE buffer at 300 V for 70-90 min. The RNA was transferred to a Zeta-Probe GT membrane (Bio-Rad) at 20V overnight at 4L °C in 0.5 X TBE. Next, the RNA was crosslinked to the membrane by UV irradiation. RiboRuler Low Range RNA ladder (Thermo Fisher Scientific) was marked by UV-shadowing. Oligonucleotide probes (Table S5) for the different RNAs were labeled with [γ-32P] ATP (166µCi/µl, 7,000 mCi/mmole, Enco) by incubating with 10 U of T4 polynucleotide kinase (New England Biolabs) at 37 °C for 1 hour and cleaned using G50 columns (Microspin™ G-50 Columns) before probing onto membrane. The membranes were visualized by Typhoon FLA 7000 phosphorimager (GE Healthcare)

#### Western blot analysis

Samples were grown to the desired density, an equivalent of OD_600_ = 1 was collected, pelleted, and immediately frozen in liquid nitrogen. Cells were mixed with 2 X Laemmli sample buffer (Bio-Rad) normalized to the cell density, and heated to 95 °C for 10 min. Samples were separated on a Mini-PROTEAN TGX 5%–20% Tris-Glycine gel (Bio-Rad) and transferred to a nitrocellulose membrane (Thermo Fisher Scientific) via semi-dry transfer. Membranes were blocked in 1 X PBST containing 3% milk, probed with a 1/1000 dilution of ANTI-FLAG M2-Peroxidase (HRP) (Sigma-Aldrich), and detected with an ECL Western Blotting Detection Kit (Tivan) using the ChemiDoc imaging system (Bio-Rad).

#### P1 transduction

P1 phage was used to horizontally transfer deletions of *E. coli* genes from the Keio collection ^88^ to a desired background. Briefly, the donor was grown with P1 for 2-3 hours till lysis appeared. Lysate was obtained by centrifuging and saving the supernatant. 100 μl of the lysate was incubated with 100 μl of recipient cells at 37 °C for 20 min. Phage activity was stopped by adding 100 μl of 1 M sodium citrate, and cells were recovered with 1 ml of LB at 37 °C with rotation. All cells were plated on an appropriate plate supplemented with Kanamycin. Colonies were grown, isolated, re-plated, and verified by colony PCR with the appropriate primers (Table S5).

#### Spontaneous transition from lysogenic to lytic cycle

Prophage cultures were grown overnight at 37 °C in LB. The next day the cultures were diluted into TBMM supplemented with 0.2% maltose and 0.0001% vitamin B1. At the designated time point, OD_600_ was measured and 1 ml of bacteria culture was collected and centrifuged. The supernatant was taken and PFU/ml was calculated as described above in the Plaque assay to evaluate the spontaneous transition from lysogenic to lytic cycle.

#### qPCR of intercellular DNA

Prophages were grown overnight in 2 ml LB. Cultures were diluted and grown to mid-log phase. 20 ml of each culture were exposed to 40 mJ/cm^2^ of 254 nm UV irradiation using UVP Crosslinker (Analytik Jena) and immediately placed back in the incubator to continue the growth. At specified time points, OD_600_ was measured and an equivalent of OD_600_ = 1 was collected, pelleted, and immediately frozen in liquid nitrogen. Intercellular DNA was purified using a Bacterial genomic DNA isolation kit (Norgen biotek) and analyzed in qPCR with specific oligonucleotide primers that were designed for each gene (Table S5) as follows. Equal amounts of DNA were loaded into a 96-well plate and DNA was quantified by CFX Connect Real-Time system (Bio-Rad) using iTaq Universal Sybr Green mix (Bio-Rad) according to manufacturer instructions. Serial dilutions of *E. coli* and lambda genomic DNA in known concentrations were used to generate a standard curve. CFX maestro analysis software (Bio-Rad) was used to determine the starting quantities of the DNA samples based on the standard curve. Reactions for each biological replicate were performed in technical triplicate.

#### In-vitro RNA synthesis

DNA templates for RNA synthesis were amplified from the MG1655 prophage genome using P1101 and P1102 for *dnaN*, and P1103 and P1104 (Table S5) for LPR1. RNAs were synthesized using T7 RNA polymerase (25 units, New England Biolabs) in a 50 µl reaction containing 1 X T7 RNA polymerase buffer, 10 mM DTT, 20 units of recombinant RNase inhibitor, 500 µM of each NTP, and 300 ng of T7 promoter containing template DNA at 37[°C for 2 hours, followed by 10 min at 70 °C. Next, Turbo DNase was added, and the reactions were incubated at 37 °C for 30 min. The RNA was purified by phenol-chloroform extraction and precipitated using 0.5 M NaCl, ethanol, and GlycoBlue. Concentration was measured with a NanoDrop (Thermo Fisher Scientific).

#### Sanger sequencing for *dnaN*

The *dnaN* sequence was PCR amplified from the bacterial genome using primers P1105 and P1106 (Table S5), digested by SacI and pstI, and cloned into pGEM-3 digested by the same enzymes. Plasmid was denatured by incubating with 0.2 mM NaOH at RT for 5 min followed by precipitation with ammonium acetate. DNA was sequenced using Sequenase version 2.0 DNA polymerase (TermoFisher scientific). Briefly, Plasmid was incubated with labeled reverse primer P1102 (Table S5) and the appropriate buffer for 15 min at 37 °C. Four separate reactions, each with one ddNTP, were prepared with DDT and the supplied enzyme. 3.2 μl of labeled plasmid were added and incubated at 37 °C for 15 min. The reaction was stopped by adding stop solution and heating to 95 °C for 3 min.

#### Structural probing and primer extension

Structural probing reactions were carried out using DMS and RNase T1. Briefly, RNA was preheated to 70 °C for 5 min, 1[pmol of *dnaN* was incubated with 10[pmol of LPR1 at 22[°C for 20 min in their respective buffers (with 100 mM MgCl), followed by incubation with 0.1 U of RNase T1 (37 °C for 5 min) or 0.3% DMS (23 °C for 5 min). RNase T1 reaction was also carried out in the presence of 2 pmole (hexameric concentration) of purified Hfq (Purified protein was kindly provided by Gisela Storz). Reactions were stopped by phenol-chloroform extraction and precipitated using 0.3 M sodium acetate and GlycoBlue. RNA concentration was measured using a NanoDrop (Thermo Fisher Scientific). 5 ng of RNA from the DMS treatment and 7 ng of RNA from the T1 RNase treatment were incubated with a [γ-^32^P] ATP (166µCi/µl, 7,000 mCi/mmole, Enco) labeled reverse *dnaN* primer P1102 (Table S5) at 70 °C for 5 min, followed by 10 min in ice. The reactions were subjected to primer extension at 42 °C for 45 min using 1 unit of MMLV-RT (Promega) and 0.5 mM of dNTPs. Extension cDNA products were analyzed on 6% acrylamide 8 M urea-sequencing gels and visualized by Typhoon FLA 7000 phosphorimager (GE Healthcare). The four samples of the Sanger sequencing for *dnaN* were loaded to the same gel and served as a reference.

#### Scarless mutations in the phage genome

Chromosomal scarless deletions within lambda were carried out as described previously ^89^. A prophage with WT lambda in its chromosome was used to delete part, or all of the *lpr1* sequence. First, a pKD46 plasmid ^90^ was transformed into the host strain. Next, a tet-sacB cassette was PCR amplified with primers that had homologous sequence to the region of the desired deletion using primers P968 and P969 for the full deletion, and P970 and P971 for seed deletion (Table S5). The amplified sequences were transformed into the cells carrying pKD46 grown with 10 mM arabinose to induce the lambda red recombinase system, and colonies sensitive to sucrose were selected using 6% sucrose plates. Deletions were verified by colony PCR and Sanger sequencing.

#### GFP translational reporter assay

GFP reporter assays were carried out essentially as described ^23^. Overnight cultures were grown in 0.75 ml of LB media supplemented with the appropriate antibiotics in a 96-deep well plate at 37 °C with constant shaking at 250 rpm. Cells were diluted in 1 ml of fresh LB medium supplemented with the appropriate antibiotics and 1 mM IPTG in a 96-deep well plate and grown at 37 °C with constant shaking at 250 rpm for 3 hours. Cells were pelleted and resuspended in filtered 1 X PBS. Fluorescence was measured using the Cytoflex flow cytometer (Beckman Coulter). The level of regulation was determined by first subtracting the auto-fluorescence and then calculating the ratio between the fluorescence signal of a strain carrying the sRNA over-expressing plasmid and the signal of a strain carrying the control plasmid. Three biological repeats were done for every sample.

#### RIL-seq analysis

RIL-seq was carried out as previously described ^22,23^. *E. coli* K-12 MG1655 cells with a FLAG-tagged Hfq were grown to exponential phase in LB medium supplemented with 10 mM MgSO_4_, separated into two flasks, and one flask was infected by lambda phage (MOI = 5). Samples were collected from these cultures 30 and 60 min following the infection and were then subjected to RIL-seq and RNA-seq analyses as described in detail previously ^22^. The libraries were sequenced by paired-end sequencing using the HiSeq 2500 system (Illumina). Experiments were done with two biological replicates. Similar treatment was done with cells carrying untagged Hfq and samples were collected 30 min after the infection. This resulted in a total of 12 samples for each of the RIL-seq and RNA-seq datasets as detailed in Table S1. The computational part of the RIL-seq is described in detail under “Data processing”.

### QUANTIFICATION AND STATISTICAL ANALYSIS

#### Data processing

Information on *E. coli* genes, transcriptional units, promoters, and terminators was sourced from EcoCyc (version 27.5; ^61^) and supplemented by manual curation in our lab. The annotation file used for various analyses is a .gff file containing the above data of *E. coli* genes and lambda phage genes.

The RNA-seq pipeline includes the following stages: splitting the data to libraries, removal of adapters, quality filtering, mapping, and counting the reads mapped to each element in the annotation file ^22^. The process_nextseq_run.py script (https://github.com/asafpr/RNAseq_scripts; version 1.1) was run on the .fastq files of the RNA-seq experiment using a fasta file containing the *E. coli* genome and the lambda genome as two chromosomes, the .gff annotation file, and -r --cutadapt_and_map_mem_per_cpu 48000 --skip_bcl2fastq --dont_delete --allowed_mismatches 3 --gene_identifier gene_name -a AGATCGGAAGAGC -A AGATCGGAAGAGC parameters. The other parameters were default.

The RIL-seq pipeline (version 0.82) includes the RNA-seq pipeline’s stages in addition to reproducibility analysis, unifying libraries, finding chimeras, and determining the statistically significant chimeras (https://github.com/asafpr/RILseq) ^22^. To get BAM files of the alignment, process_nextseq_run.py script (version 1.1) was run on the .fastq files of the RIL-seq experiment, which were split to libraries, with the same parameters as mentioned before, except to --skip_split that was added. The script map_chimeric_fragments.py was run for each BAM file, with the parameters -r and -t with the .gff annotation file. The other parameters were default.

#### Checking reproducibility between libraries

For checking reproducibility, the RILseq_significant_regions.py script was used twice, once running with --all_interactions parameter, and then running with --only_singles in addition to the --all_interactions parameter. In both runs the files from EcoCyc with the information about the *E. coli* transcriptional units, promoters, and terminators were used with the --bc_dir parameter, the --min_odds_ratio was set to 1, the parameter --total_RNA was used with the corresponding file from the total RNA-seq pipeline results, the parameters -- total_reverse and --ribozero were used, the parameter -g was used with the genome fasta file, and the parameter --BC_chrlist were used with the appropriate names. The other parameters were defaults. In addition, the plot_regions_interactions.py script was run with default parameters.

After unifying biological replicates, the script RILseq_significant_regions.py was used to find the S-chimeras with the parameters mentioned above, but without --all_interactions and --only_singles parameters, and with --refseq_dir parameter to get the genes descriptions.

#### DESeq2

Differential gene expression analysis between two different conditions was conducted by DESeq2 (version 1.36.0) ^34^. DESeq2 analysis provided the log (fold change) and the p-value of the change (corrected by multiple hypothesis testing). DESeqDataSet was created by DESeqDataSetFromMatrix with default parameters, the results were re-ordered by the relevel function and then were used by the function DESeq with default parameters. Finally, the DESeq outputs were sent to the result function and were written to files.

#### Circos plots

Circos plots were created by Rcircos library (version 1.2.2) ^91^ in R (version 4.2.0). A data.frame with information about the *E. coli* and the lambda genomes was created and loaded to RCircos.Set.Core.Components function. The Circos parameters were received by RCircos.Get.Plot.Parameters function, and the chrom.paddings, base.per.unit and text.size parameters were set to 700, 45.137051, and 0.4, respectively. Those parameters were reset by RCircos.Reset.Plot.Parameters function. The Circos plots were plotted by RCircos.Set.Plot.Area, RCircos.Draw.Chromosome.Ideogram, and RCircos.Label.Chromosome.Names functions. Files with the locations and the corresponding numbers of the scale marks were created, and loaded by the function RCircos.Gene.Connector.Plot, and the function RCircos.Gene.Name.Plot. Finally, files that represent the chimeras’ locations and colors were created and loaded by the function RCircos.Link.Plot with the parameter lineWidth that was set to a vector full of 0.00001 (for the Circos plots with all the interactions) or 2 (for the Circos plots with the interactions of LPR1, LPR2, and 6S RNA) where its length is the number of chimeras.

#### Motif discovery

To find motifs in the sequences sited in chimeras with LPR1, each chimera was extended with 30 bases from each side, and overlapping sequences were concatenated, to avoid biases in the results. The motifs were searched in the resulting sequences by MEME tool (version 5.4.1) ^92,93^ with -rna -minw 6 -maxw 15 -nmotifs 3 -mod zoops parameters, the other parameters were default.

To find whether the motif that was found with MEME is complementary to LPR1 sequence, MAST (version 5.4.1) ^58^ was run on the reverse complement sequence of LPR1 with the MEME output, and the default parameters (-ev 10 -mt 0.0001).

#### Venn diagrams

Venn diagrams of S-chimeras between two different groups are generated with the venn2 function of the python (version 3.9) package matplotlib-venn (version 0.11.7) ^94^.

#### Bar plots

Bar plots that represent the percentage of S-chimeras were generated using the python (version 3.9) package matplotlib (version 3.5.1) ^94^.

#### Heatmaps

Heatmaps of the chimeras’ annotations were created with the heatmap function of the python (version 3.9) package seaborn (version 0.11.2) ^95^.

#### Interaction networks

RNA-RNA interaction networks were generated using Cytoscape (version 3.9.1) ^96^. json file that contains the data about the genes and the connections between them was generated and uploaded to Cytoscape.

#### 3D folding and interaction model

AlphaFold 3 server ^69^ was used to compute the folding and interaction of PreS, *dnaN*, and Hfq. The input was PreS and *dnaN* RNA sequences and 6 copies of the Hfq protein sequence. The output .cif file was loaded to UCSF ChimeraX, a software developed by the Resource for Biocomputing, Visualization, and Informatics at UCSF ^97^, to color it and optimize its display. Hfq C-tails were removed from the display of the final 3D model. The Hfq*-dnaN*-PreS complex video was generated by using the ChimeraX export option and was edited using CapCut ^98^.

#### LPR1 conservation

To find whether LPR1 is conserved in other genomes, we submitted its sequence to BLAST search ^99^ (version 2.12.0) against the Nucleotide database with the command blastn and -db nt -remote -max_target_seqs 100000 parameters. Next, for the bacteria and the phages separately, we generated a phylogenetic tree based on one representative genome of each species that was found by BLAST, using BV-BRC ^100^ Bacterial Genome Tree tool with 5 Max Allowed Deletions and Duplications. Finally, we downloaded the bacteria and phage trees, joined them to one tree, and loaded it to MEGA11 (version 11.0.13) ^101^. To visualize the conservation between lambda and the other genomes, for each representative genome, we extract the region around the sequence that was found with BLAST. Next, Easyfig (version 2.2.5) ^102^ was used with these extracted regions and the lambda genome to generate the images.

#### Figure preparation

Figures and the Supplementary Figures were created using BioRender (BioRender.com).

## Acknowledgments

We extend our gratitude to Susan Gottesman for providing the WT lambda strain, Shoshy Altuvia for the lambda strains CI-and CIII864, and Sivan Pearl-Mizrahi for the *E. coli* harboring the CI857::kan prophage. We thank Gisela Storz for work conducted by S.M. in her lab and the NICHD Molecular Genomics Core for sequencing the RNA-seq and RIL-seq libraries. We also thank Maya Elgrably-Weiss and Shoshy Altuvia for their assistance with the structural probing experiments. We are grateful to Hongen Zhang, Hanah Margalit and her lab members for assisting with the computational analyses, and to Alon Shainer-Melamed for helping with the video editing. We are grateful to the Melamed lab for their insightful discussions and thank the Melamed lab, Gisela Storz, Hanah Maraglit, and Reut Shainer for their valuable comments on the manuscript. Additionally, we appreciate Rachel Marianovsky for her help with lab maintenance tasks. This work utilized the computational resources of the Hebrew University Research Computing Services. This work was supported by the Israel Science Foundation (Grants 826/22 and 2859/22).

## Author contributions

S.M. conceived the project. S.M., R.N., A.S., and T.N. designed, performed, and analyzed the experiments. R.B. performed all the computational analyses. S.M, A.S, and R.B. prepared the figures and wrote the manuscript. S.M. supervised the project.

## Declaration of generative AI and AI-assisted technologies in the writing process

During the preparation of this work the author(s) used ChatGPT and Perplexity in order to improve language and readability. After using this tool/service, the author(s) reviewed and edited the content as needed and take(s) full responsibility for the content of the publication.

