## Supplementary Figures and data for "Phage-encoded small RNA hijacks host replication machinery to support the phage lytic cycle"

### SUPPLEMENTAL INFORMATION

#### Table of Contents

**Supp. Figure S1. *E. coli* transcriptome and sRNA interactome during lambda infection, and data reproducibility** (related to Figure 1)

**Supp. Figure S2. Lambda-encoded sRNAs expression and RIL-seq target sets** (related to Figure 2)

**Supp. Figure S3. *lpr1* mutants' expression and phenotypes, and regulation of *alaS*** (related to Figures 3 and 4)

**Supp. Figure S4. *dnaN* structural probing using T1 RNase** (related to Figure 5)

**Supp. Figure S5. Conservation of LPR1 among bacteria and phages** (related to Figure 6)

**Supp. Figure S6. Confidence of 3D modeling of Hfq-*dnaN*-PreS complex** (related to Figure 7)

**Supp. Table legends**

#### SUPPLEMENTARY FIGURES

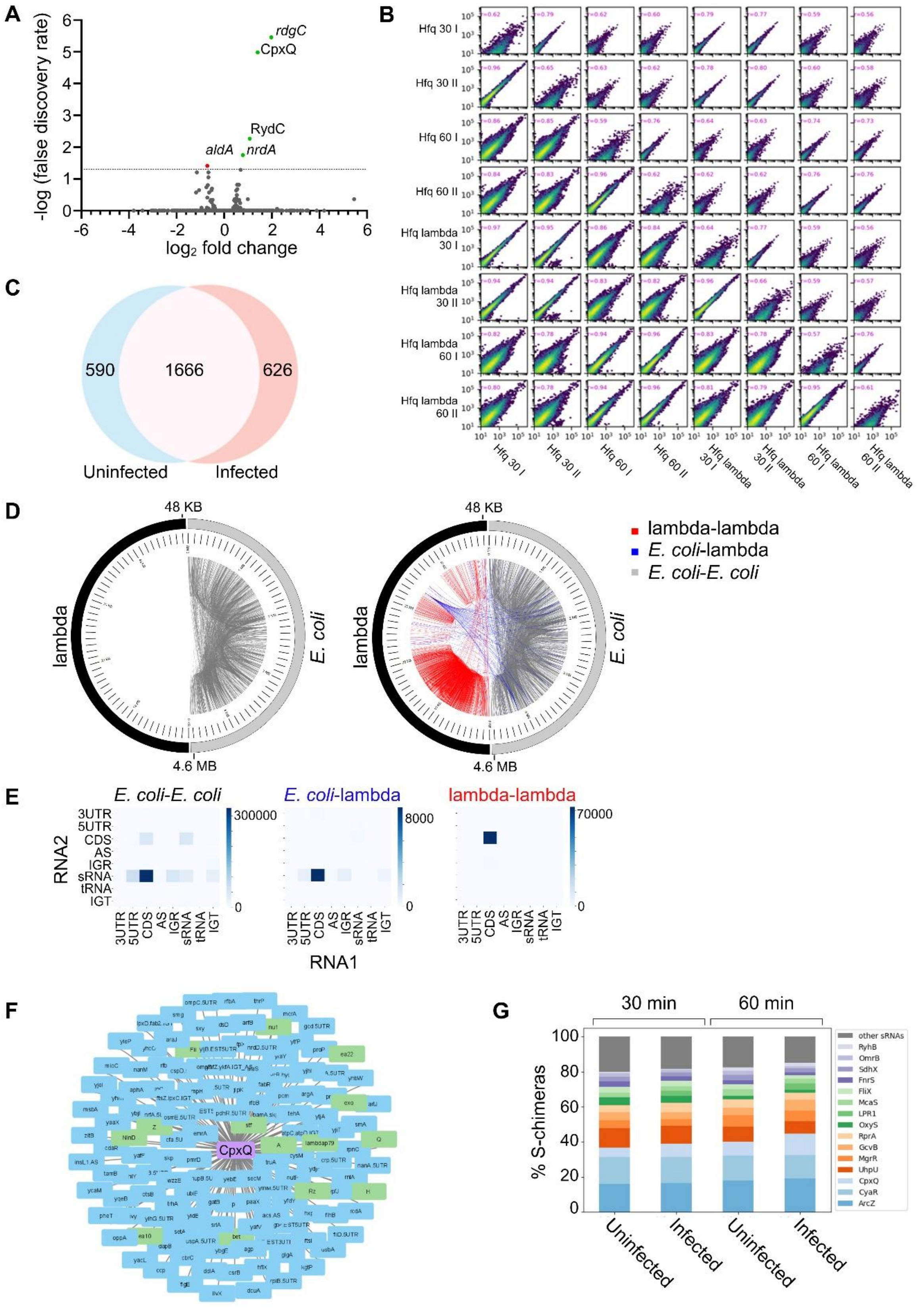

**Supp. Figure S1. *E. coli*-lambda transcriptome and sRNA interactome during lambda infection, and data reproducibility** (related to Figure 1)

**(A)** Change in gene expression in *E. coli* 30 min following lambda infection is represented by a volcano plot. Genes with a p-adjusted value $< 0.1$ (above the dashed bar) are considered as exhibiting statistically significant changes in expression. Genes that show statistically significantly increased expression are colored green and genes that show statistically significantly decreased expression are colored red. Genes that do not statistically significantly change are colored gray. Prophage genes were removed since it was not possible to determine if the mapped reads belonged to the prophage genes or the homologous lambda genes. See more information in Table S2.

**(B)** The reproducibility of results within same-condition libraries was assessed for single fragments and chimeric fragments. Scatter plots were used to compare the sequenced fragments between the two libraries. Each point represents the number of fragments mapped to a 100-nucleotide region in both libraries. Dot intensity ranges from blue (fewer fragments) to red (more fragments). Plots below the diagonal show results for single fragments while plots above the diagonal display results for chimeric fragments. Diagonal plots represent specific libraries, comparing single and chimeric fragments mapped to the same genome region. Library names are as listed in Table S1.

**(C)** Overlap of *E. coli* chimeras between infected and uninfected bacteria. Venn diagram represents the overlap between the *E. coli* chimeras at 30 min after the infection and the chimeras obtained in *E. coli* without infection. The location of RNAs in the chimeras as RNA1 and RNA2 was ignored in this analysis.

**(D)** Circos plots of *E. coli* and lambda chimeras in uninfected (left) and infected samples (right) 30 min after the infection. Half of each circle represents the *E. coli* genome, and the other half represents the lambda genome. Each edge in a circle represents a chimera between two regions of the genomes. *E. coli*-*E. coli* chimeras, *E. coli*-lambda chimeras and lambda-lambda chimeras colored by gray, blue, and red, respectively. Each scale mark represents 100,000 bases in the *E. coli* half and 1,000 bases in the lambda half.

**(E)** Total number of chimeric fragments for each combination of genomic elements in *E. coli* - *E. coli* chimeras, *E. coli*–lambda chimeras, and lambda–lambda chimeras, 30 min after infection. Rows represent the first RNA in the chimera, and columns represent the second RNA in the chimera.

**(F)** CpxQ interaction network 30 min after infection. *E. coli*-encoded genes are labeled in blue and lambda-encoded genes are labeled in green. The network was drawn by Cytoscape [1].

**(G)** Distribution of Hfq-bound sRNAs forming S-chimeras at 30 and 60 min post-infection in infected and uninfected datasets.

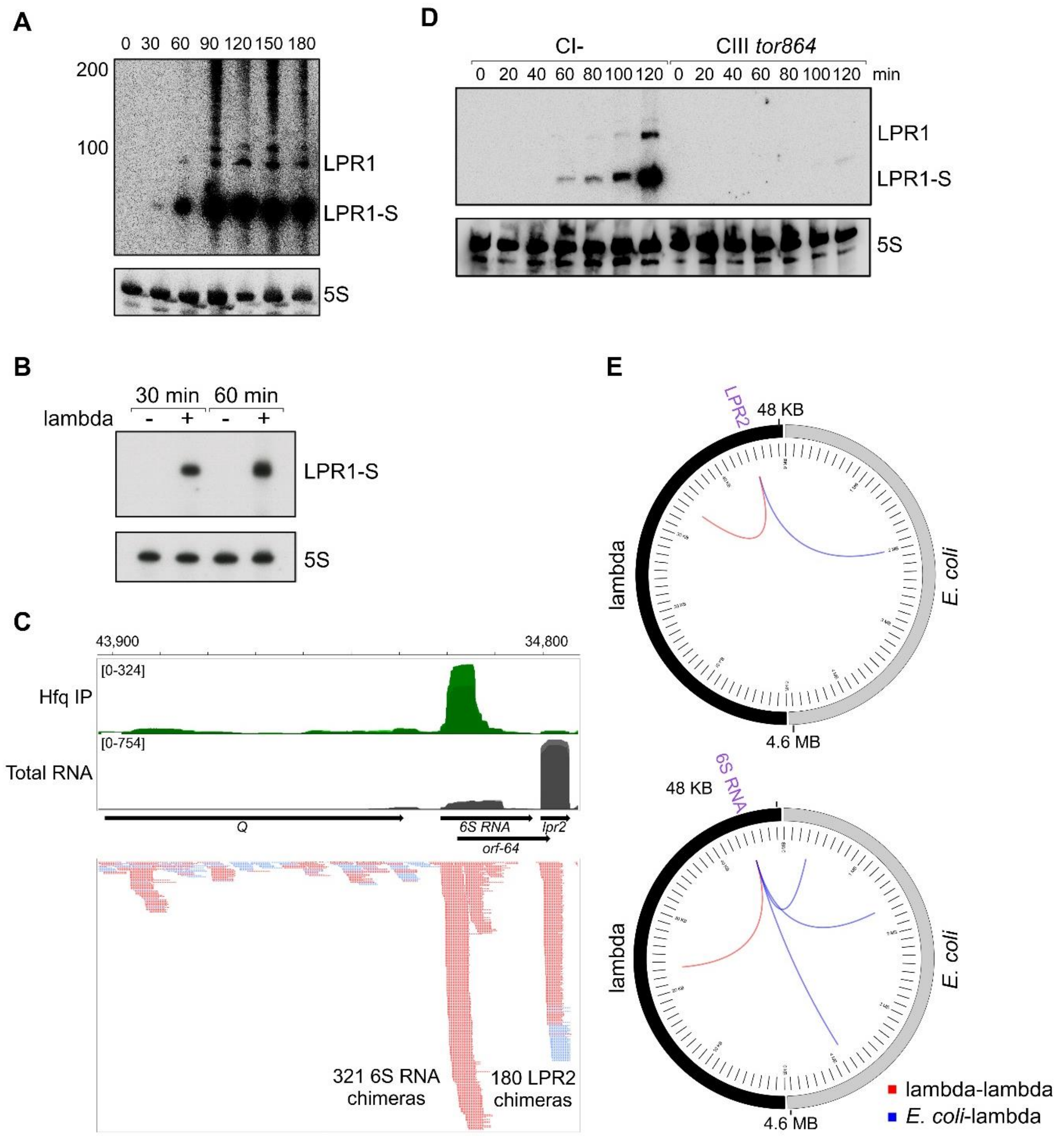


**Supp. Figure S2. Lambda-encoded sRNAs expression and RIL-seq target sets** (related to Figure 2)

**(A)** Northern blot analysis documenting LPR1 levels throughout infection of *E. coli* with WT lambda phage (MOI = 0.0015). Longer exposure of the membrane used in Figure 2C detects LPR1 starting from 30 min post-infection. The 5S RNA served as a loading control and the same 5S panel used in Figure 2C.

**(B)** Northern blot analysis using the samples used for the RNA-seq experiment described in Table S1. LPR1 is detected 30- and 60- min following infection of *E. coli* with WT lambda phage (MOI = 5). The 5S RNA served as a loading control.

**(C)** Browser image of *lpr2 and 6S RNA* region. Top: Hfq IP (green) and total RNA (gray). Normalized read count ranges are shown in the upper left. Bottom: chimeras of LPR2 and 6S RNA. The red and blue colors indicate if LPR2 and 6S RNA were found as RNA1 or RNA2 in the chimera, respectively. Data shown is of 30 min after the infection.

**(D)** Northern blot analysis showing LPR1 is expressed when *E. coli* is infected with lambda strain CI- but not with CIII *tor864*. Cultures were grown to mid-log and then infected with lambda strain CI- or CIII tor864. Samples were collected every 20 min for 2 h. The 5S RNA served as a loading control.

**(E)** Representation of LPR2 (top) and 6S RNA (bottom) chimeras, 30 min after infection. Circos plots were drawn as described in Figure 1C.

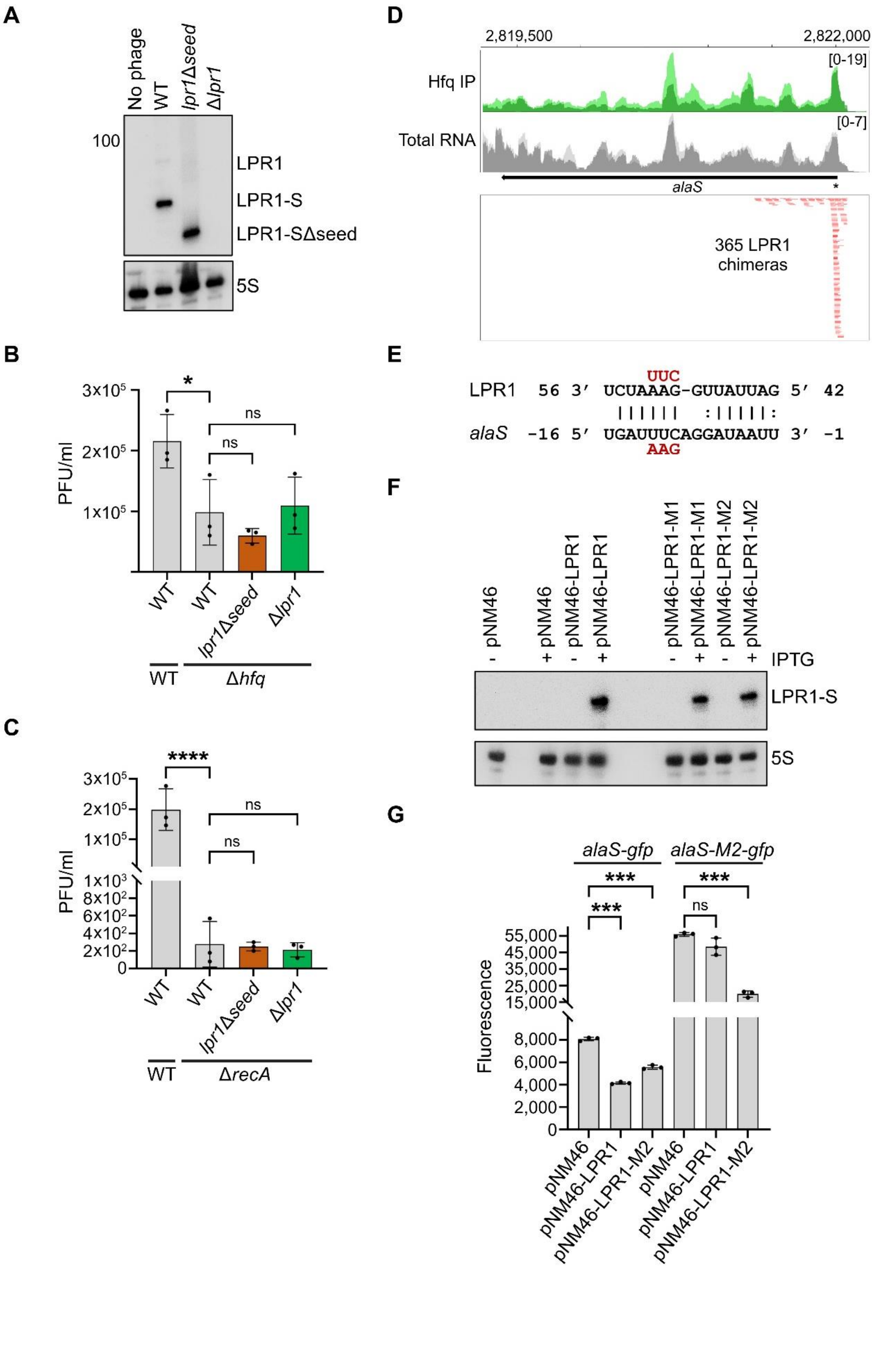
A
No phage
WT
lpr1Δseed
Δlpr1
100
LPR1
LPR1-S
LPR1-SΔseed
5S
B
3x10^5
2x10^5
1x10^5
PFU/ml
*
ns
ns
WT
WT
lpr1Δseed
Δlpr1
WT
Δhfq
C
****
3x10^5
2x10^5
1x10^5
1x10^3
8x10^2
6x10^2
4x10^2
2x10^2
0
PFU/ml
ns
ns
WT
WT
lpr1Δseed
Δlpr1
WT
ΔrecA
D
2,819,500
2,822,000
Hfq IP
[0-19]
Total RNA
[0-7]
alaS
*
365 LPR1 chimeras
E
UUC
LPR1 56 3′ UCUAAAG-GUUAUUAG 5′ 42
alaS -16 5′ UGAUUUCAGGAUAAUU 3′ -1
AAG
F
pNM46
pNM46
pNM46-LPR1
pNM46-LPR1
pNM46-LPR1-M1
pNM46-LPR1-M1
pNM46-LPR1-M2
pNM46-LPR1-M2
- + - + - + - +
IPTG
LPR1-S
5S
G
alaS-gfp
alaS-M2-gfp
***
***
***
ns
55,000
45,000
35,000
25,000
15,000
8,000
6,000
4,000
2,000
0
Fluorescence
pNM46
pNM46-LPR1
pNM46-LPR1-M2
pNM46
pNM46-LPR1
pNM46-LPR1-M2

**Supp. Figure S3. *lpr1* mutants' expression and phenotypes, and regulation of *alaS*** (related to Figures 3 and 4)

**(A)** Northern blot analysis documenting LPR1 levels with activation of lambda from prophages (WT, *lpr1*Δ*seed*, Δ*lpr1*) by UV irradiation. Bands are detected only in WT and *lpr1*Δ*seed* lambda strains. The 5S RNA served as a loading control.

**(B)** Similar transition of WT lambda and mutated prophages to the lytic life cycle in Δ*hfq* background. PFU/ml was detected from a spontaneous lysate of WT and mutated prophages in Δ*hfq* background. Lambda WT strain in WT *E. coli* background serves as a reference. The two mutants do not show a decrease in the progeny of phages upon spontaneous induction in comparison to WT lambda. Results are the average of 3 biological replicates. Error bars represent one SD.

**(C)** Similar transition of WT lambda and mutated prophages to the lytic life cycle in Δ*recA* background. PFU/ml was detected from a spontaneous lysate of WT and mutated prophages in Δ*recA* background. Lambda WT strain in WT *E. coli* background serves as a reference. The two mutants do not show a decrease in the progeny of phages upon spontaneous induction in comparison to WT lambda. Results are the average of 3 biological replicates. Error bars represent one SD.

**(D)** Browser image of *alaS* region. Top: Hfq IP (green) and total RNA (gray). Normalized read count ranges are shown in the upper right. Bottom: chimeras of *alaS* with LPR1. The red color indicates that *alaS* was found as RNA1 in the chimera. The asterisk indicates the position of the base pairing with LPR1. Data shown is of 30 min after the infection.

**(E)** Base pairing between *alaS* and LPR1 with sequences of mutants assayed (colored in red). Numbering is from AUG of *alaS* mRNA and +1 of LPR1 sRNA.

**(F)** Northern blot analysis documenting comparable levels of LPR1 and its mutants, expressed from the pNM46 vector upon addition of 1 mM IPTG. The 5S RNA served as a loading control. **(G)** LPR1 reduces *alaS*-*gfp* reporter fusion based on reporter assays of *alaS*-*gfp* expressed from pXG10-SF. This reduction is not observed when using LPR1-M2. Compensatory mutation in alaS-*gfp* restores the downregulation by LPR1. Average fluorescent values are based on three biological replicates. Error bars represent one SD. One-way ANOVA comparison was performed to calculate the statistical significance of the change in plaque formation in (B) and (C) in GFP signal in (G) (ns = not significant, *=p < 0.05, ***=p < 0.0001, ****=p < 0.0001).

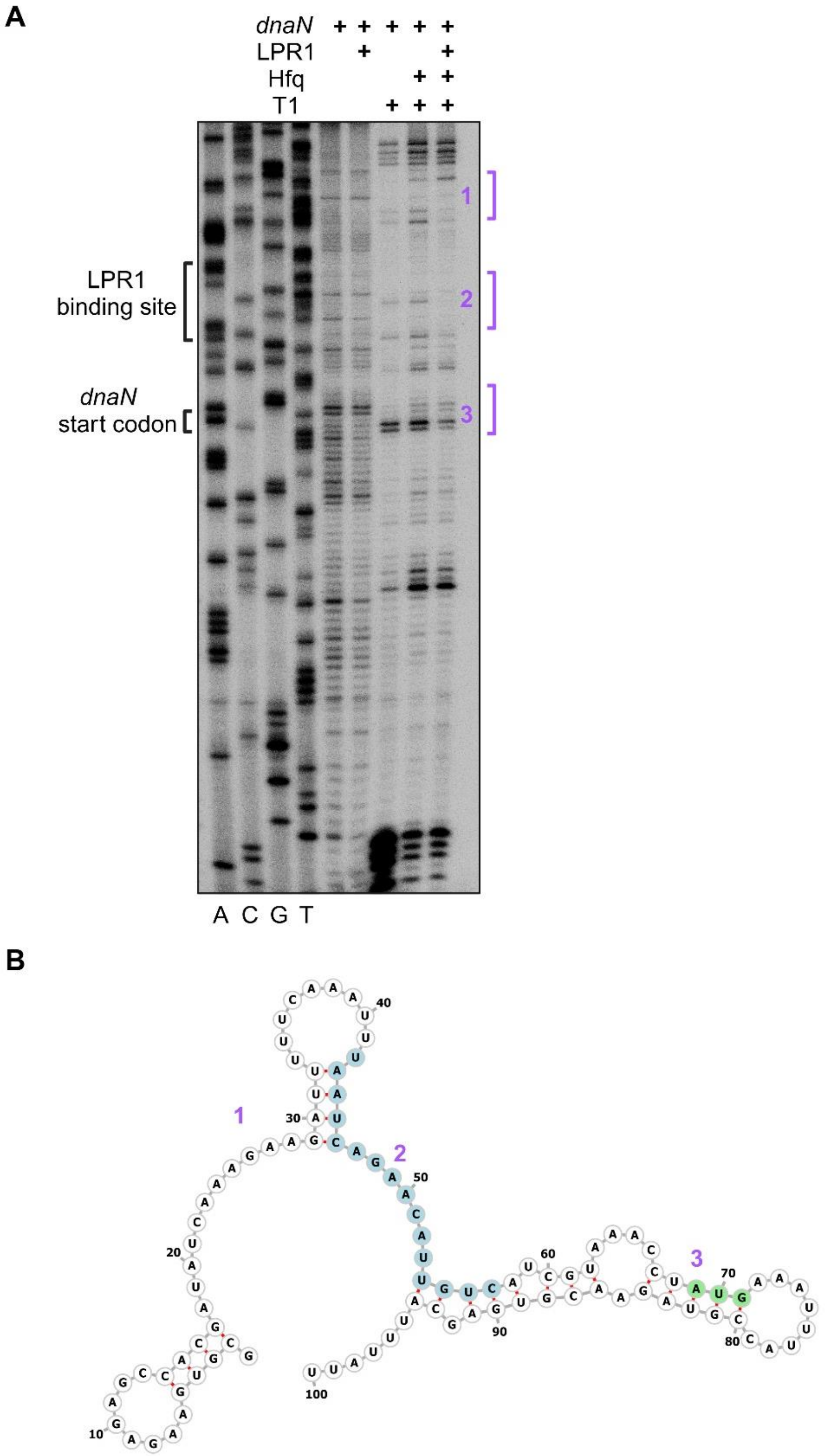


**Supp. Figure S4.** ***dnaN*** **structural probing using T1 RNase** (related to Figure 5)

**(A)** Structural probing of *dnaN* with and without LPR1 treated with RNase T1. RNase T1 cleaves RNA at 3′-end of guanosine. Regions where the binding of LPR1 changes the T1 RNase cleavage pattern are numbered (1-3), while region 2 overlaps LPR1 binding site as shown in Figure 4B. The left 4 lanes are sanger sequencing of the *dnaN* sequence used in this assay, ACTG represents the ddNTP that was added. Note that the observed band pattern represents the *dnaN* complementary sequence. Lanes 5 and 6 are untreated *dnaN* without- and with - LPR1, respectively. Lane 7 is *dnaN* treated with RNase T1. Lane 8 is *dnaN* treated with RNase T1 in the presence of Hfq. Lane 9 is *dnaN* treated with RNase T1 in the presence of Hfq and LPR1. In all lanes, a radioactive Primer (P1102) was used for the primer extension.

**(B**) Secondary structure of the *dnaN* RNA used for the structural probing assay. Drawn using forna [2]. The first 100 nt of the *dnaN* RNA used are shown. LPR1 binding site is highlighted in blue, *dnaN* start codon is highlighted in green, and positions changed in the structural probing assay are labeled with purple numbers.

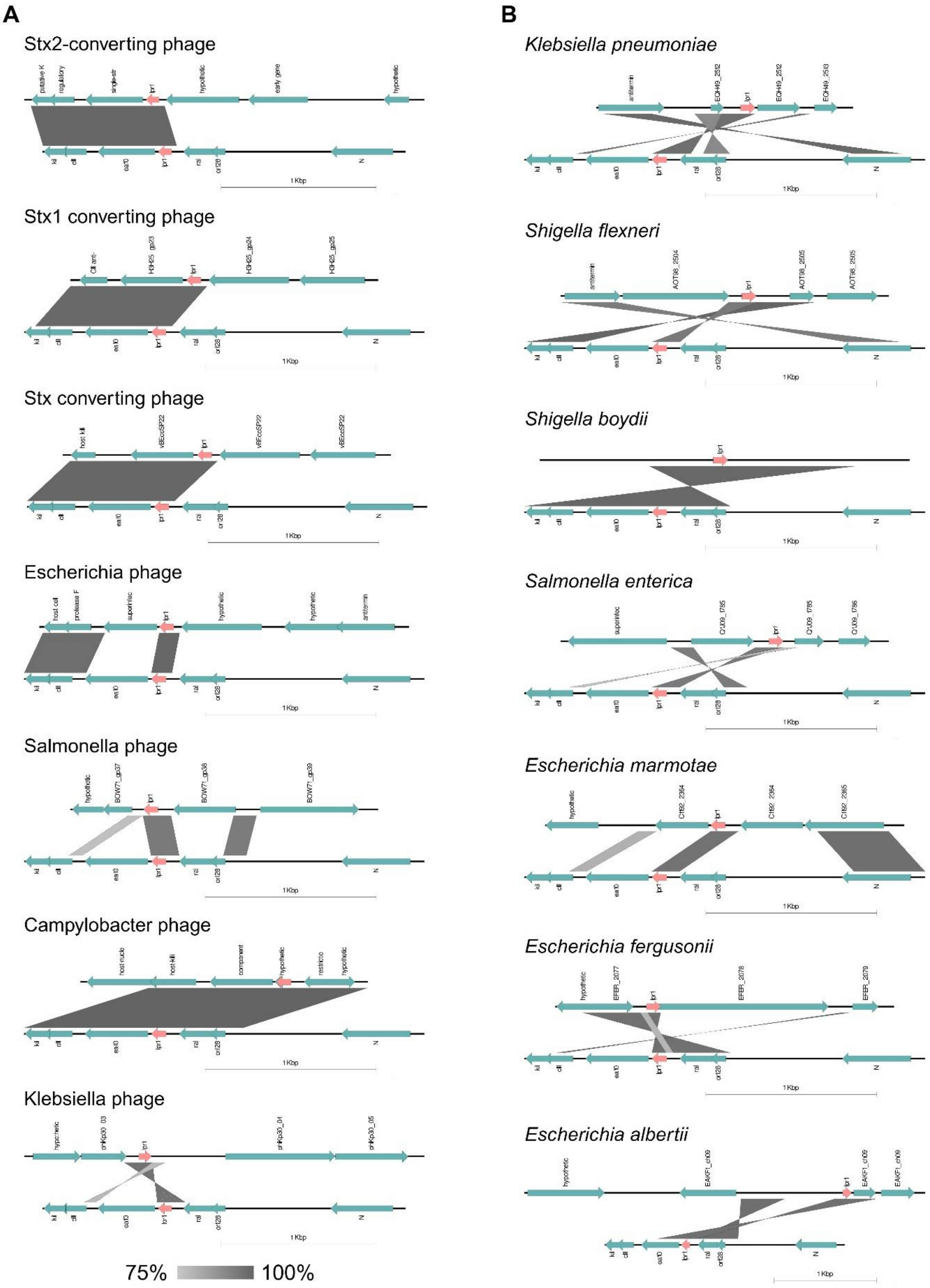
A
Stx2-converting phage
Stx1 converting phage
Stx converting phage
Escherichia phage
Salmonella phage
Campylobacter phage
Klebsiella phage
75%
100%
B
Klebsiella pneumoniae
Shigella flexneri
Shigella boydii
Salmonella enterica
Escherichia marmotae
Escherichia fergusonii
Escherichia albertii

**Supp. Figure S5. Conservation of LPR1 among bacteria and phages** (related to Figure 6) Representation of the conservation of LPR1's region in selected bacterial species **(A)** and phages **(B)**. The similarity between regions on the genomes is marked by gray. The scale represents the identity percentage according to BLAST. The sequences' full names are listed in Table S4.

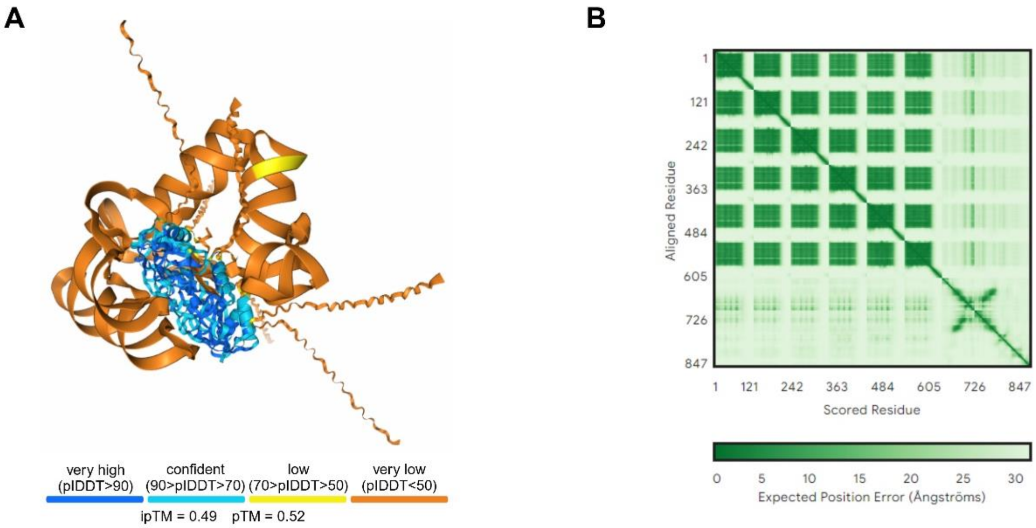


**Supp. Figure S6. Confidence of 3D modeling of Hfq-*dnaN*-PreS complex** (related to Figure 7)

**(A)** 3D model of the Hfq-dnaN-PreS complex. Prediction of the 3D structure of the Hfq-dnaN-PreS complex by AlphaFold 3 [3] recapitulates our experimental results for dnaN-PreS interaction. The color scale for plDDT (predicted local-distance difference test) represents the confidence measure to estimate the accuracy of the structure predictions.

**(B)** Assessment of the quality of protein and RNA structure predictions. The plot represents the predicted position error for each residue (amino acid for Hfq and nucleotide for the RNAs) along the x-axis, assuming alignment of the predicted and true structures based on the residue shown on the y-axis..

#### SUPP. TABLE LEGENDS

**Table S1. Number of fragments in deep sequencing libraries (related to Figures 1, 2, 3, S1, and S2).** The table describes the different libraries used in all experiments with statistics regarding the number of sequenced fragments. The total number of fragments includes the number of fragments available after splitting the libraries according to the barcodes. For the RIL-seq libraries, the RIL-seq computational pipeline [4] was used to evaluate the results of each library separately as well as the unified libraries per condition.

**Table S2. RNA levels in different RNA-seq datasets (related to Figures 1A and S1A).** Total RNA library reads were subject to differential expression analyses conducted with DESeq2 [5].

Fold Change: Fold change between the groups.

log2 Fold Change: log2 fold change between the groups.

P-value: Wald test p-value.

padj: Benjamini-Hochberg adjusted p-value.

-LOG(pADJ): -log of Benjamini-Hochberg adjusted p-value.

baseMean: The average of the normalized counts taken over all samples.

lfcSE: standard error of the log2 Fold Change estimate.

stat: Wald statistic.

The “prophage gene analysis” tab highlights the challenges of mapping reads to prophage genes that have homologs in lambda. This analysis displays mapped reads for ten prophage genes and their homologous lambda genes, with or without lambda infection, using mismatch-free mapping. For certain prophage genes (*rzpD*, *nohA*, *tfaD*, and *ybcW*), in all replicates of infected samples, more reads were mapped to the prophage gene when mapping to the *E. coli* genome alone, compared to mapping to both *E. coli* and lambda genomes. Almost all the reads that were mapped to these prophage genes when mapping to lambda and *E. coli* genomes, were mapped to the homolog gene when mapping to lambda genome alone. In addition, over 40% of reads mapped to the prophage gene in the *E. coli* genome were reassigned to the homologous lambda gene when both genomes were used. Since the mapping was done without mismatches, this suggests that for some prophage genes, it is difficult to discern whether mapped reads belong to them or to the homologous lambda genes.

### reads mapped to lambda, mapping to lambda and *E. coli*: number of reads that were mapped to the lambda gene when mapping to the *E. coli* and lambda genomes.
### reads mapped to prophage, mapping to lambda and *E. coli*: number of reads that were mapped to the prophage gene when mapping to the *E. coli* and lambda genomes.
### reads mapped to prophage, mapping to *E. coli*: number of reads that were mapped to the prophage gene when mapping to the *E. coli* genome.
### reads mapped to lambda when mapping lambda and *E. coli*, but mapped to prophage when mapping to *E. coli* alone: number of reads that were mapped to the lambda genes when mapping to lambda and *E. coli* genomes but mapped to the prophage genes when mapping to *E. coli* genome alone.
% reads that mapped to lambda when mapping to lambda and *E. coli*, from reads that mapped to prophage when mapping to *E. coli* alone: fraction of reads from "# reads mapped to lambda when mapping lambda and *E. coli*, but mapped to prophage when mapping to *E. coli* alone"

column out of all the reads that were mapped to the prophage genes when mapping to *E. coli* genome alone.

### reads mapped to prophage when mapping to *E. coli* and lambda, but mapped to lambda when mapping to lambda alone: number of reads that were mapped to prophage genes when mapping to *E. coli* and lambda but were mapped to the lambda genes when mapping to lambda genome alone.

**Table S3. Statistically significant chimeric fragments in the RIL-seq experiment (related to Figures 1, 2, 3, S1, and S2).** RIL-seq RNA pairs identified in unified datasets. The table includes all interactions between two RNAs, which were supported by statistically significant chimeras. A pair of RNAs might appear more than once if it involves multiple interacting regions or if it appears in the chimera once as RNA1-RNA2 and once as RNA2-RNA1. Coordinates are based on the genome of *E. coli* K12 MG1655 (NC_000913.3) and lambda genome (J02459.1). The table includes data from BioCyCTM pathway/genome database under license from SRI International.

Name: Common name of the gene (additional information is included in the EcoCyc ID column).

### of chimeric fragments: Number of chimeras supporting the interaction.

### of libraries: Number of individual libraries where this interaction was revealed as statistically significant. "U" denotes an interaction that was identified only when in a unified library.

Description: Description of the gene products taken from RefSeq.

Chromosome: Defines if the RNA was mapped to *E. coli* or lambda.

Normalized Odds Ratio: Odds Ratio multiplied by the relative enrichments of RNA1 and RNA2 on Hfq.

Odds Ratio: (K/L)/(M/N), where K= Number of chimeric fragments of RNA1-RNA2, L=number of other fragments involving RNA2, M=number of other fragments involving RNA1, N=number of all other fragments (that do not involve RNA1 and RNA2).

Fisher's exact test p-value: p-value for observing at least this number of chimeric fragments given their background frequencies on Hfq. The Odds Ratio provides the effect size of the test.

Genomic annotation: e.g. sRNA, CDS, 3´UTR, etc.

RNA1 from: Position of the first nt of the most 5´ chimera mapped to the first RNA.

RNA1 to: Position of the first nt of the most 3´ chimera mapped to the first RNA.

Strand: The genome strand the sequence was mapped to.

RNA2 from: Position of the last nt of the most 5´ chimera mapped to the second RNA.

RNA2 to: Position of the last nt of the most 3´ chimera mapped to the second RNA.

Other fragments of RNA1: Number of fragments in which the first RNA appears as first, including single fragments.

Other fragments of RNA2: Number of fragments in which the second RNA appears as second, including single fragments.

Total other fragments: Number of fragments in the experiment excluding the above.

RNA1 in total RNA (# of reads): The sum of the number of fragments of RNA1 in total RNA libraries.

RNA2 in total RNA (# of reads): The sum of the number of fragments of RNA2 in total RNA libraries.

lib norm IP RNA1: RNA1 normalized number of reads in RIL-seq library lib norm IP RNA2: RNA2 normalized number of reads in RIL-seq library

lib norm total RNA1: RNA1 normalized number of reads in total RNA library

lib norm total RNA2: RNA2 normalized number of reads in total RNA library

RNA1 IP/total ratio: (fraction of RNA1 fragments in RIL-seq library) / (fraction of total RNA of RNA1 in total RNA library).

RNA2 IP/total ratio: (fraction of RNA2 fragments in RIL-seq library) / (fraction of total RNA of RNA2 in total RNA library).

EcoCyc ID: The accession number of the gene in the EcoCyc database. When the RNA was mapped to a region outside a CDS, the name is followed by 5UTR or 3UTR in case it resides in an annotated UTR (in EcoCyc), EST5UTR or EST3UTR if the UTR is unknown and the interaction is 100 nt upstream or downstream the CDS (or shorter if these regions spanned another transcript or were more likely to be a UTR of the neighboring transcript). Two gene names and IGR or IGT represent a binding region located between two genes in two different transcription units (IGR) or on the same transcription unit (IGT). AS stands for RNA mapped to the antisense of a gene.

* The table is sorted according to the # of chimeric fragments.

**Table S4. Conservation of PreS in bacteria and phages (related to Figures 6 and S5).**

The table describes the conservation of PreS in different bacteria and phages according to the BLAST results and lists the genomes that were used to create the phylogenetic tree and the conservation figures.

species: species that PreS's sequence was found in them by BLAST.

number: number of different genomes from each species that was found by BLAST.

representative genome: the genomes that were used to create the phylogenetic tree and the conservation figures.

genome id: the GenBank number of the representative genome.

Notes: notes about the representative genome or the species.

**Table S5. List of strains, plasmids, and oligonucleotides used in this work (Related to Methods).**
